# Low dose Cold Atmospheric Plasma induces membrane oxidation, stimulates endocytosis and enhances uptake of nanomaterials in Glioblastoma multiforme cells

**DOI:** 10.1101/805192

**Authors:** Zhonglei He, Kangze Liu, Laurence Scally, Eline Manaloto, Sebnem Gunes, Sing Wei Ng, Marcus Maher, Brijesh Tiwari, Hugh J Byrne, Paula Bourke, Furong Tian, Patrick J Cullen, James F Curtin

## Abstract

Cold atmospheric plasma (CAP) has demonstrated synergistic cytotoxic effects with nanoparticles, especially promoting the uptake and accumulation of nanoparticles inside cells. However, the mechanisms driving the effects need to be explored. In this study, we investigate the enhanced uptake of theranostic nanomaterials by CAP. Numerical modelling of the uptake of gold nanoparticle into U373MG Glioblastoma multiforme (GBM) cells predicts that CAP may introduce a new uptake route. We demonstrate that cell membrane repair pathways play the main role in this stimulated new uptake route, following non-toxic doses of dielectric barrier discharge CAP (30 s, 75 kV). CAP treatment induces cellular membrane damage, mainly via lipid peroxidation as a result of reactive oxygen species (ROS) generation. Membranes rich in peroxidated lipids are then trafficked into cells via membrane repairing endocytosis. We confirm that the enhanced uptake of nanomaterials is clathrin-dependent using chemical inhibitors and silencing of gene expression. Therefore, CAP-stimulated membrane repair increases endocytosis and accelerates the uptake of gold nanoparticles into U373MG cells after CAP treatment. Our data demonstrate the utility of CAP to model membrane oxidative damage in cells and characterise a previously unreported mechanism of membrane repair to trigger nanomaterial uptake which will be useful for developing more efficient deliveries of nanoparticles and pharmaceuticals into cancer cells for tumour therapy and diagnosis. This mechanism of RONS-induced endocytosis will also be of relevance to other cancer therapies that induce an increase in extracellular RONS.

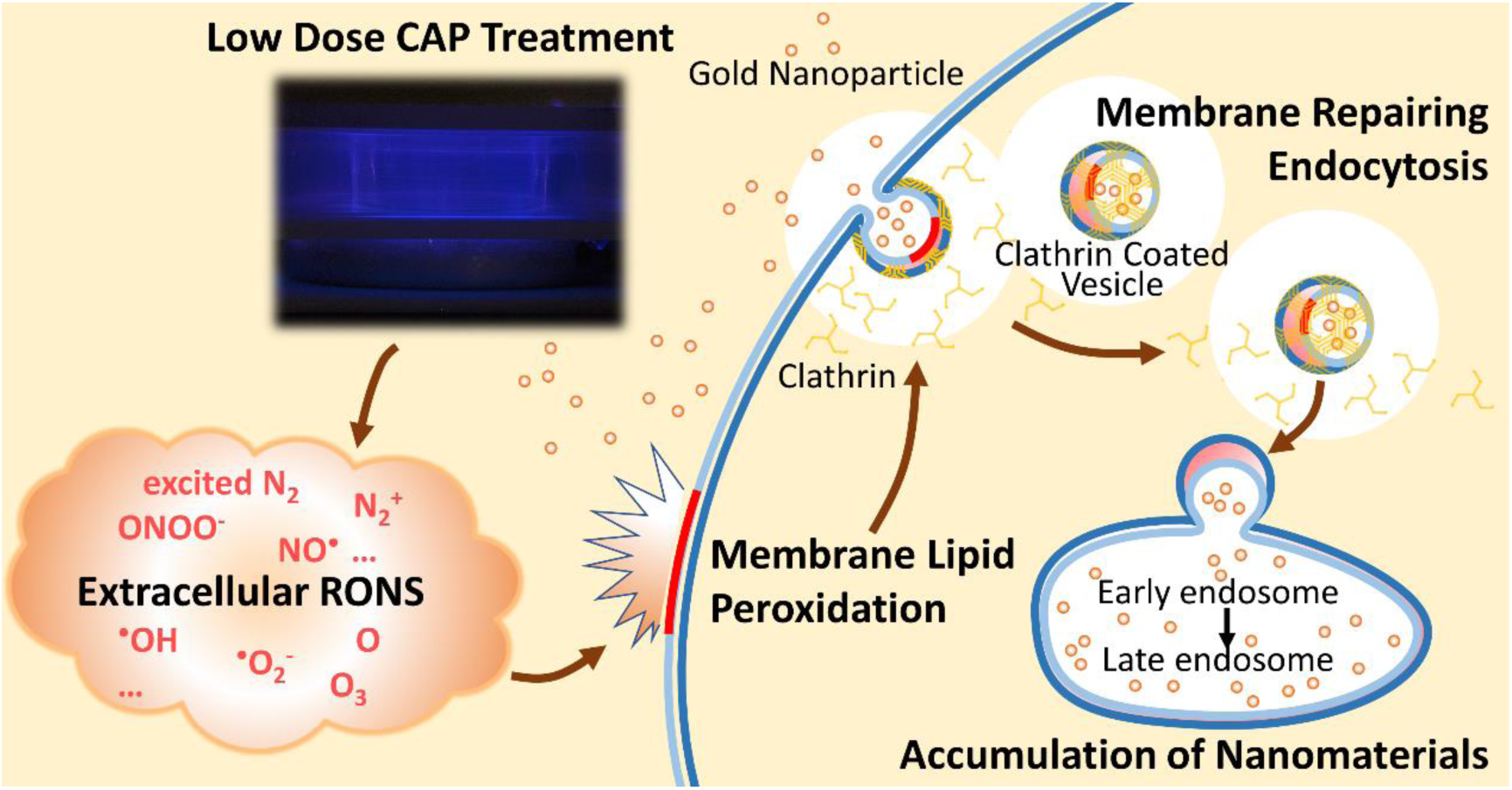

## Introduction

Cold atmospheric plasma (CAP) is increasingly studied for applications across the food industry, medicine, energy storage and for driving catalytic reactions. Technological developments and preclinical studies have led to CAP testing in a growing number of clinical trials for cancer treatment^1, 2^. Research is ongoing to explore the combination of CAP with other cancer therapies, including nanotechnology-based, radio- and chemotherapy^3–5^.

Gold nanoparticles (AuNPs) are considered to be weakly or non-toxic to human cells and highly effective for delivery through the blood brain barrier, especially for ∼20 nm diameter AuNPs^6–8^. Meanwhile, AuNPs are known to be readily manufactured and designed for targeting delivery of various compounds into cells. Therefore, they have emerged as a promising reagent, combined with CAP, for anti-cancer therapy in recent studies^4, 9, 10^. In our previous study, we explored the potential of a combination treatment of CAP with gold nanoparticles, which showed promising synergistic cytotoxicity to U373MG Glioblastoma multiforme (GBM) cells^11^. The accelerated uptake and accumulation of AuNPs in U373MG cells induced by CAP can enhance the efficiency of pharmaceuticals delivery for tumour treatment and diagnosis.

In general, the citrate-capped cationic AuNPs may absorb serum proteins onto their surface in cell culture medium and thereby stimulate receptor-mediated endocytosis pathways, including clathrin-mediated, caveolae-mediated and clathrin/caveolae independent endocytosis^8^. Without special surface functionalisation, AuNPs enter cells and are trapped in vesicles^8, 12, 13^ or enter the nucleus, depending on their size/shape^14, 15^. It has also been demonstrated that clathrin-mediated endocytosis is the specific mechanism of normal AuNPs cellular uptake^16^. Meanwhile, AuNPs with functionalised surface chemistries/ligands can directly penetrate the membrane and enter the cytoplasm^17^. However, the detailed mechanism whereby CAP and AuNPs have synergistic biological effects on cancer cells and the uptake of AuNPs affected by CAP needs to be further explored.

CAP generates a unique physical and chemical environment, including generating short- and long-lived reactive nitrogen species (RNS, e.g. excited N_2_, N_2_^+^, ONOO^−^ and NO^•^, etc.) and reactive oxygen species (ROS, e.g. ^•^OH, O, ^•^O_2_^−^ and O_3_, etc.), photons as well as heat, pressure gradients, charged particles, and electrostatic and electromagnetic fields, many of which are known to induce biological effects^18–20^. Parallels to this can be found in phagocytes of the immune system. Enzymatic production of reactive oxygen and nitrogen species (RONS) along with various hypohalous acids, especially hypochlorites, play a significant role in respiratory bursts, also known as oxidative bursts, which are used in the clearance of tumour cells by phagocytic immune cells including neutrophils, macrophages and monocytes^21^. Anti-cancer cytotoxicity induced by respiratory bursts has been shown to induce spontaneous regression in mouse tumour models^22–24^. ROS has emerged as a double-edged sword to cancer cells. Evidence shows that higher levels of ROS are generated in cancer cells by comparison with normal cells, which is attributed to the higher metabolic activities and more rapid proliferation of transformed cells^25^. Hence, the cellular antioxidant system works under a heavier load to protect tumour cells from oxidative stress, suggesting it may be possible to selectively overload and eliminate them with locally induced ROS production^26^.

Reactive species can induce a free radical chain reaction in the membrane lipids, known as lipid peroxidation/oxidation, which leads to oxidative degradation of the lipids and therefore a disruption of the membrane function and induced injury and disorder to cells. The peroxidated lipid products can induce further propagation of the free radical reactions^27^. It has been shown that several ROS and RNS generated by CAP as well as natural biological processes can induce cell injures via lipid peroxidation^28^. For example, as an oxidant prominent in air pollution, O_3_ has been proven to be responsible for the lipid peroxidation damage in lung cells^29–31^. Hydroxyl radicals (^•^OH) react with various cellular components, including membrane lipid^32^ Superoxide (^•^O ^−^) can form peroxynitrite (ONOO^−^), which is able to initiate lipid peroxidation, after reacting with nitric oxide (NO)^32^. RNS, such as NO_2_ and ONOOH, also interact with lipids to form nitrated lipids, which have been demonstrated to play roles in vascular and inflammatory cellular signalling pathways^28^.

The following study of CAP-accelerated AuNPs uptake was carried out using a high voltage dielectric barrier discharge (DBD) contained reactor which has been previously described and characterised^11^. U373MG cells were treated with a low dose of CAP treatment previously demonstrated to be non-lethal, at a voltage output of 75 kV for 30 s^33^. Numerical modelling of the uptake of AuNPs, indicates that the CAP treatment stimulated a new uptake route, which can be clathrin-mediated membrane repairing rapid endocytosis according to experimentally observed behaviour. RONS generation by CAP is characterised using H_2_DCFDA, optical emission spectroscopy and quantitative colorimetric titration methods. We provide evidence, using the TBARS assay and a C11-BODIPY lipid peroxidation sensor, that CAP generated RONS induce lipid peroxidation, membrane damage and thereby activate the membrane reparation via rapid endocytosis. We employ various clathrin and caveolin specific inhibitors and clathrin silencing to further determine that the CAP-induced endocytosis of AuNP and membrane damage response is clathrin-dependent.

## Results

### Numerical modelling of the uptake of AuNPs by GBM cells

The accumulation of AuNPs inside U373MG cells was monitored using atomic absorption spectroscopy and the dose response curve of AuNPs with or without CAP treatment has been presented in previous study ^11^. We further analysed the data according to a simulated uptake model to better understand the possible mechanism of CAP-stimulated AuNP uptake. The uptake of nanoparticles by cell populations *in vitro* has previously been modelled according to a phenomenological rate equation approach^34–36^, and the approach can be extended to further investigate the role of CAP in AuNP uptake by U373MG Glioma Cells.

The rate of uptake of AuNPs into a cell can be described by the equation:

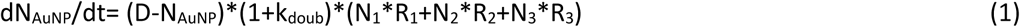

where N_AuNP_ is the number of internalised gold nanoparticles, D is the initial dose of AuNPs, (D-N_AuNP_) allows for the depletion of the applied AuNP dose, and k_doub_ is the doubling time of the cells. N_1_/R_1_, N_2_/R_2_and N_3_/R_3_ allow for three different principle uptake pathways, with respective limiting capacities of N and rates R. The first two terms describe independent active and passive uptake mechanisms, respectively, with limiting cellular capacities N_1_(0) = N_1max_ and N_2_(0) = N_2max_, such that:

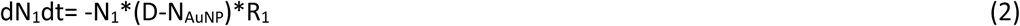

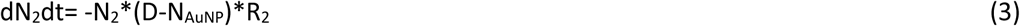

Figure 1 (solid blue line) shows the simulated uptake of AuNPs, normalised to the maximum uptake observed for AuNP + CAP, for the case of R_1_ = 3 x 10^-3^ hr^-1^, R_2_ = 2.5 x 10^-5^ hr^-1^, R_3_ =0, which faithfully reproduces the experimentally observed behaviour. Quenching of the active uptake of AuNPs by NaN_3_ is best simulated by addition of a further term in equation 2, such that

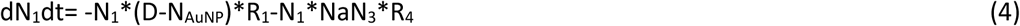

where NaN_3_ is the effective dose of sodium azide, and R_4_ allows for the rapid depletion of the active uptake pathway. The experimentally observed uptake was well simulated (solid orange line) by a value of R_4_ = 3 x 10^-5^ hr^-1^, keeping all other rates as before.

In simulating the increased uptake of AuNPs upon CAP treatment, it was noted that the enhancement of a single pathway described by equations (2-4) by CAP treatment, by increasing a single uptake rate, did not faithfully reproduce the experimentally observed behaviour, as the uptakes were limited by the parameters N_1max_ and N_2max_. Rather, faithful reproduction of the observed behaviour required the introduction of independent uptake mechanisms for untreated and CAP treated AuNP uptake, an observation which was critical to the interpretation of the effects of CAP treatment on the cells. Such a pathway can be represented by:

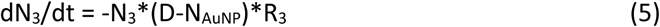

such that N_3_(0) = N_3max_. Upon the application of CAP, the enhanced uptake was well fitted (solid yellow line) by R_3_ = 2.5 x 10^-4^ hr^-1^, keeping all other rates as before.

The modelling process therefore indicates that CAP treatment increases the capacity of the U373MG cells to uptake AuNPs by introducing a new uptake channel. The modelling parameters employed are detailed in Table 1. Note that the parameters relating to the limiting cell uptake, N_nmax_, were determined by the definition of the dose as 100 mg/mL. Furthermore, the process was one of simulation, rather than a mathematical fitting, so the parameters should be considered within ∼10% confidence.

**Figure 1.**
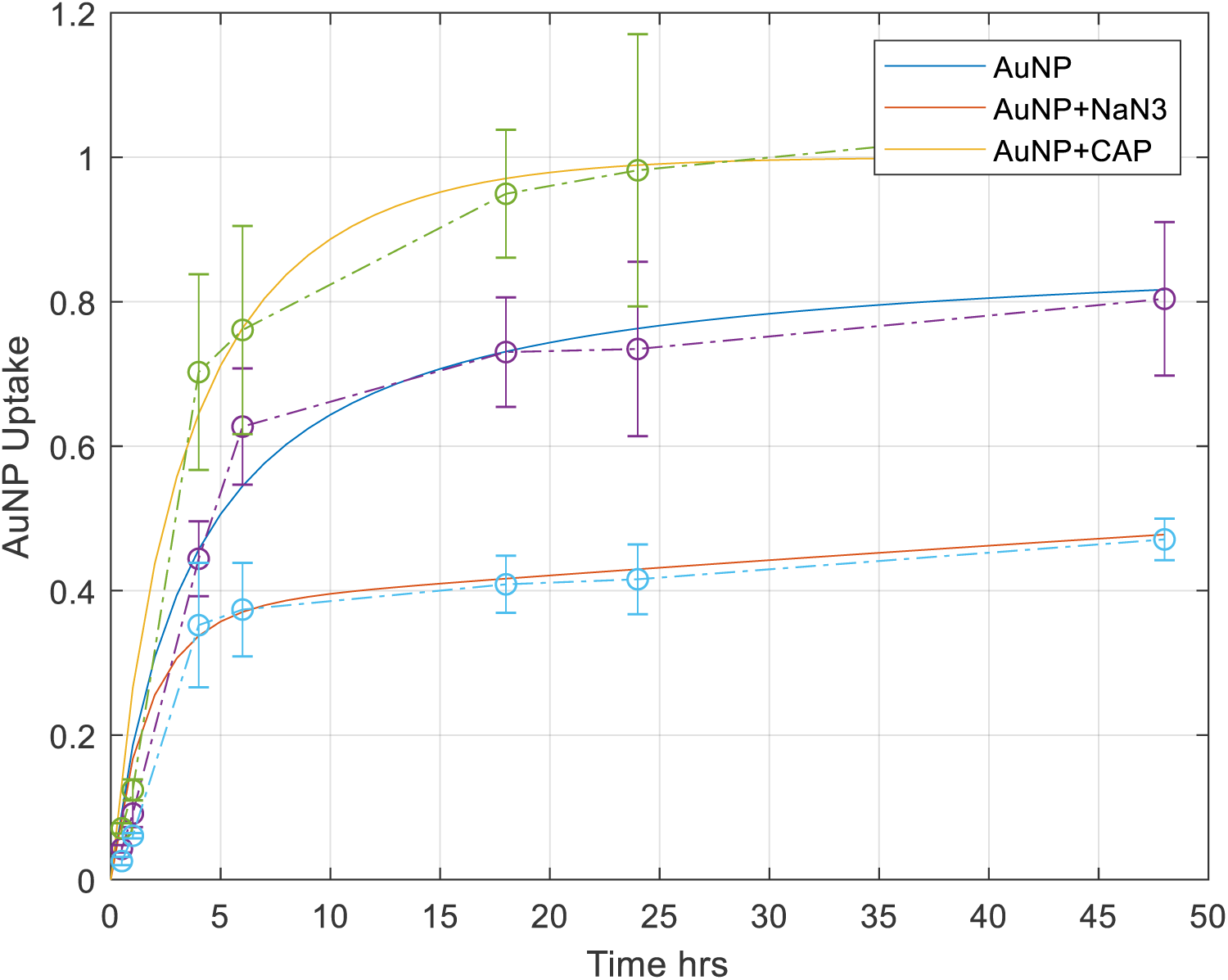
Modelling uptake of AuNPs. Numerical modelling of experimental data from our previous study^11^ (shown with open circles) was carried out for AuNP uptake (blue solid line), AuNP uptake quenched by incubation of the cells with NaN_3_ (orange solid line), AuNP uptake on application of low dose CAP (yellow solid line).

**Table 1.** The modelling parameters employed in Figure 1.

| | $N_{1max}$ | $N_{2max}$ | $N_{3max}$ | $\text{NaN}_3$ | $R_1 \text{ hr}^{-1}$ | $R_2 \text{ hr}^{-1}$ | $R_3 \text{ hr}^{-1}$ | $R_4 \text{ hr}^{-1}$ |
| --- | --- | --- | --- | --- | --- | --- | --- | --- |
| **AuNP** | 75 | 150 | 400 | 8500 | $3 \times 10^{-3}$ | $2.5 \times 10^{-5}$ | 0 | 0 |
| **AuNP + $\text{NaN}_3$** | 75 | 150 | 400 | 8500 | $3 \times 10^{-3}$ | $2.5 \times 10^{-5}$ | 0 | $3 \times 10^{-5}$ |
| **AuNP + CAP** | 75 | 150 | 400 | 8500 | $3 \times 10^{-3}$ | $2.5 \times 10^{-5}$ | $2.5 \times 10^{-4}$ | 0 |

### Reactive species generated by CAP treatment

To further identify the possible uptake pathways affected by CAP treatment and the pathways implicated by the numerical modelling of our data, ROS generation of 30 s, 75 kV CAP treatment was first investigated. Using optical emission spectroscopy (OES) and Gastec gas detector tubes, several reactive species were measured. These included N_2_, N ^+^, ^•^OH, and O_3_. OES emission intensities from the N_2_ second positive system (SPS) were measured at 315, 337, 357, and 377 nm, the N_2_ ^+^ first negative system (FNS) at 391 nm, and ^•^OH at 310 nm were measured every 7.5 s during 30 s CAP treatment, with the first point taken just when the CAP was ignited. The data demonstrated a relatively constant ROS and RNS production throughout the 30 s treatment (Figure 2a).

**Figure 2.**
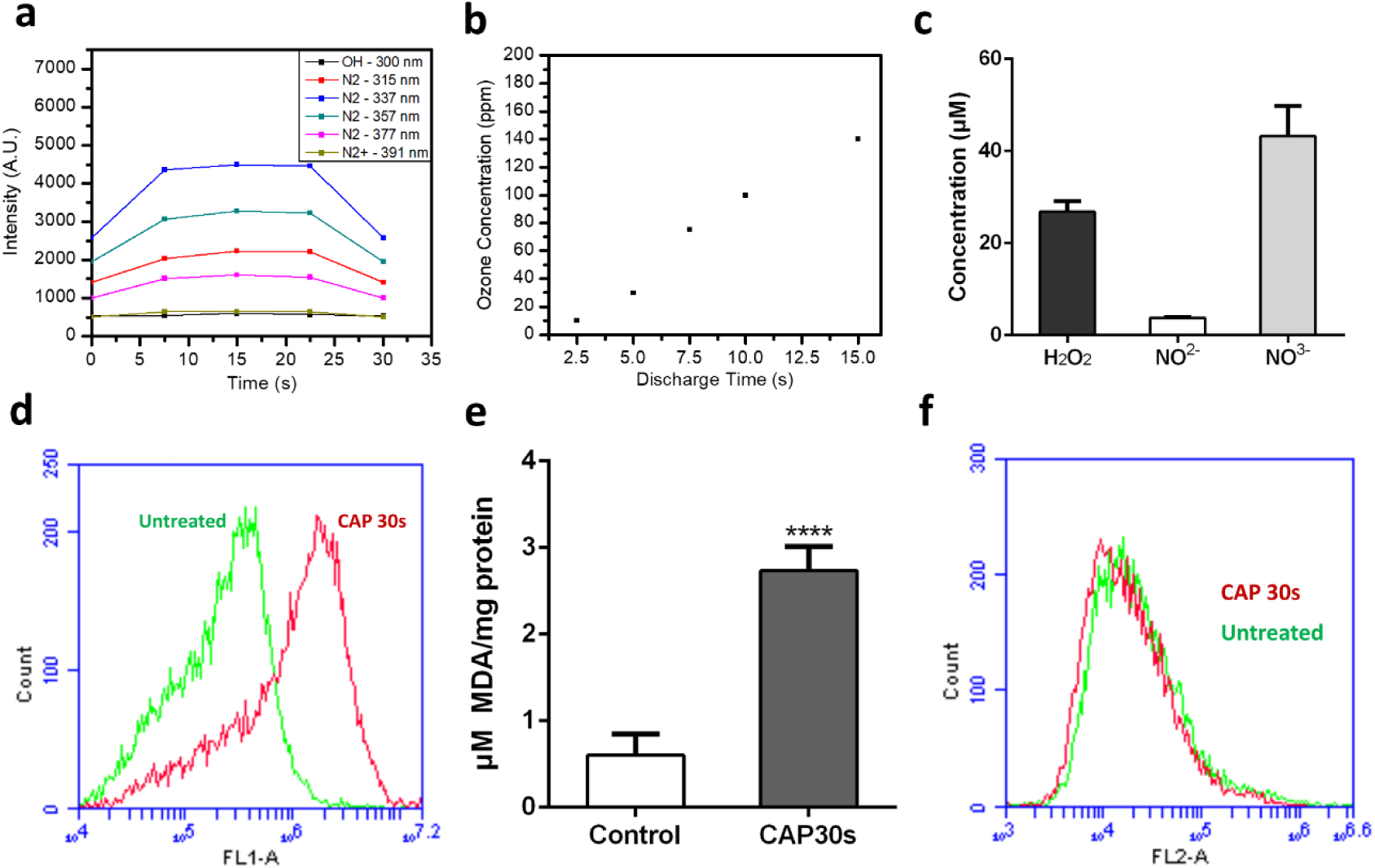
Measurement of reactive species generated by CAP treatment by OES and H_2_DCFDA, TBARS assay and PI staining. (**a**) Emission intensities of excited N_2_ molecules, N_2_ ^+^ and ^•^OH. (**b**) Concentration of O_3_ measured during CAP treatment. (**c**) The concentrations of hydrogen peroxide, nitrite and nitrate were measured in CAP-treated culture medium. (**d**) Fluorescence level of intracellular oxidised H_2_DCFDA was measured via Flow cytometry, left curve (green, untreated cells), right curve (red, CAP-treated cells). (**e**) U373MG cells were incubated for 24 hours after CAP treatment (0-30 s, 75 kV) and then collected and analysed to detect cellular MDA level using TABARS assay. (**f**) 30 mins after 30 s of 75 kV CAP treatment, cells were stained with PI for 5 min, then measured

The electron energy distribution function (EEDF) of the plasma was also determined. As seen in Supplementary Figure S1, the EEDF remained close to a ratio of 7 during CAP treatment, which indicated that the electron energies were distributed more so on the lower end of the energy scale (11 - 12 eV) than the higher energy levels (18.8 eV). The low variability of the EEDF indicated that the electric field was stable, and that the formation of the reactive species was in a steady state manner. Using Gastec ozone detector tubes, the concentrations of generated O_3_ in the extracted gas were measured post-discharge of CAP treatment (Figure 2b), revealing significantly increasing levels of O_3_ recorded over the discharge time, which became saturated at the maximum labelled value of O_3_ detection tube after 15 s. The high level of ozone generation may give explanation why no detectable or low emission of NO, O, NO_x_ (NO_2_, NO_3_, N_2_O_2_, N_2_O_3_, and N_2_O_4_), ^•^OH and N ^+^ measured in air using OES^37^(As seen in Supplementary Discussion).

Having confirmed RONS generation in plasma, the concentration of hydrogen peroxide (H_2_O_2_), nitrite (NO^−^_2_) and nitrate (NO^−^_3_) in culture media was next measured. As seen in Figure 2c, the RONS generated in CAP-treated phenol red-free medium are presented after normalising to the values of untreated medium. By direct comparison previous results^38^, CAP treatment of culture media for 30 s generated very low amounts of H_2_O_2_ (∼20 μM), NO^2-^(∼5 μM), and NO^3-^ (∼30 μM). These concentrations are at least 15-fold and 200-fold lower than the IC_50_ values we measured previously for U373MG cells (315 µM, >1200 µM and >600 µM respectively) and therefore are essentially non-toxic^39^.

To further investigate CAP induced ROS generation in U373MG cells, H_2_DCFDA, a cell permeable ROS fluorescence sensor, was preloaded into cells before CAP treatment for 0.5 h. After CAP treatment, cells were collected and the fluorescence of H_2_DCFDA was measured using flow cytometry. As seen in Figure 2d, CAP treatment induced increased intracellular H_2_DCFDA fluorescence, compared to the untreated group. The mean fluorescence was observed to significantly increase by 4-5 fold above untreated controls (Supplementary Figure S1).

### CAP treatment induces lipid peroxidation

ROS induces lipid peroxidation of the cell membrane^40, 41^. The TBARS assay detects the peroxidised lipid by-product, Malondialdehyde (MDA), as an indicator of the level of lipid peroxidation. After 30 s of CAP treatment, U373MG cells were incubated for 24 hours. The level of the lipid peroxidation indicator malondialdehyde (MDA) was significantly higher in CAP treated cells compared with the control group (Figure 2e), which demonstrated a high level of lipid peroxidation induced by CAP treatment in U373MG cells.

In our previous study, U373MG cells showed a high resistance to CAP-induced cytotoxicity compared with Hela cells. The IC_50_ of CAP treatment (75 kV) was determined to be 74.26 s (95% confidence range of 47.24–116.8 s) for U373MG cells^33^. We have previously determined that 60 s CAP treatment induces rapid permeabilisation of U373MG cell membranes^42^. Figure 2f demonstrated no significant increase of PI uptake following 30 s CAP treatment compared to the control, which indicated that the permeability of the cell membrane to PI, an indicator of membrane integrity, was not significantly affected by 30 s CAP treatment and CAP-induce lipid peroxidation. This is in agreement with our previous findings, where 30 s CAP treatment induces very low, non-significant levels of toxicity in U373MG cells 48 hours post treatment (18.52%, SEM=5.41%)^11^.

CAP induced lipid peroxidation was further analysed and visualised using the lipid peroxidation fluorescent sensor C11-BODIPY (581/591) with flow cytometry and confocal microscopy. BODIPY fatty acid is a lipophilic fluorescent dye which contains a polyunsaturated butadienyl portion that can be oxidised, leading to a shift of the emission peak from around 590 nm (Orange) to around 510 nm (Green). It has been demonstrated that C11-BODIPY reacts with various ROS and RNS and shows constant shifting of the emission peak^43^. As seen in Figure 3a, the confocal imaging showed that there was a significantly stronger green fluorescence emission in CAP-treated cells compared to the untreated cells. Furthermore, C11-BODIPY can be used as tracers of lipid trafficking, especially trafficking of oxidised membrane in this case. Confocal images demonstrated green fluorescent vesicle-like structures which could be oxidised lipid membrane fragments within endosomes or lysosomes (Figure 3a). To determine the level of lipid peroxidation, more than 60 cells were analysed using ImageJ software for each group. The data showed a significant increase (∼2 fold) of green fluorescent intensity in the CAP-treated group (Figure 3b) and the orange fluorescent intensity, which presented that the non-oxidised C11-BODIPY decreased for the CAP-treated group compared to the control group (Supplementary Figure S2). Furthermore, the fluorescence levels of C11-BODIPY in the two groups were confirmed with flow cytometry. As seen in Figure 3c, 30s CAP-treated cells showed a significantly increased signal in channel FL1-A, thereby affirming a significant increase in lipid peroxidation induced by CAP treatment (30 s, 75 kV). To calculate integrated density from strongly fluorescent vesicle-like areas only, confocal images were converted to 8-bit form, then inverted and adjusted with same threshold using ImageJ. The integrated density of those highlighted areas was then measured and analysed. As seen in Supplementary Figure S5, the total integrated density of CAP-treated group was significantly higher than untreated control (∼3 fold, ****p<0.0001). Furthermore, due to the limitation of fluorescence channels, the LysoTracker™ Deep Red, which is highly selective for acidic organelles, was used to determine the location of oxidised form of C11-BODIPY together with oxidised membrane fragments. Supplementary Figure S3 showed the co-staining of oxidised C11-BODIPY (green) and LysoTracker™ Deep Red (red). The white arrows pointed out examples of vesicle-like structure stained with C11-BODIPY which was co-localised with acidic organelles (such as lysosomes, late endosomes, etc.) as seen in Supplementary Figure S3, middle& right.

**Figure 3.**
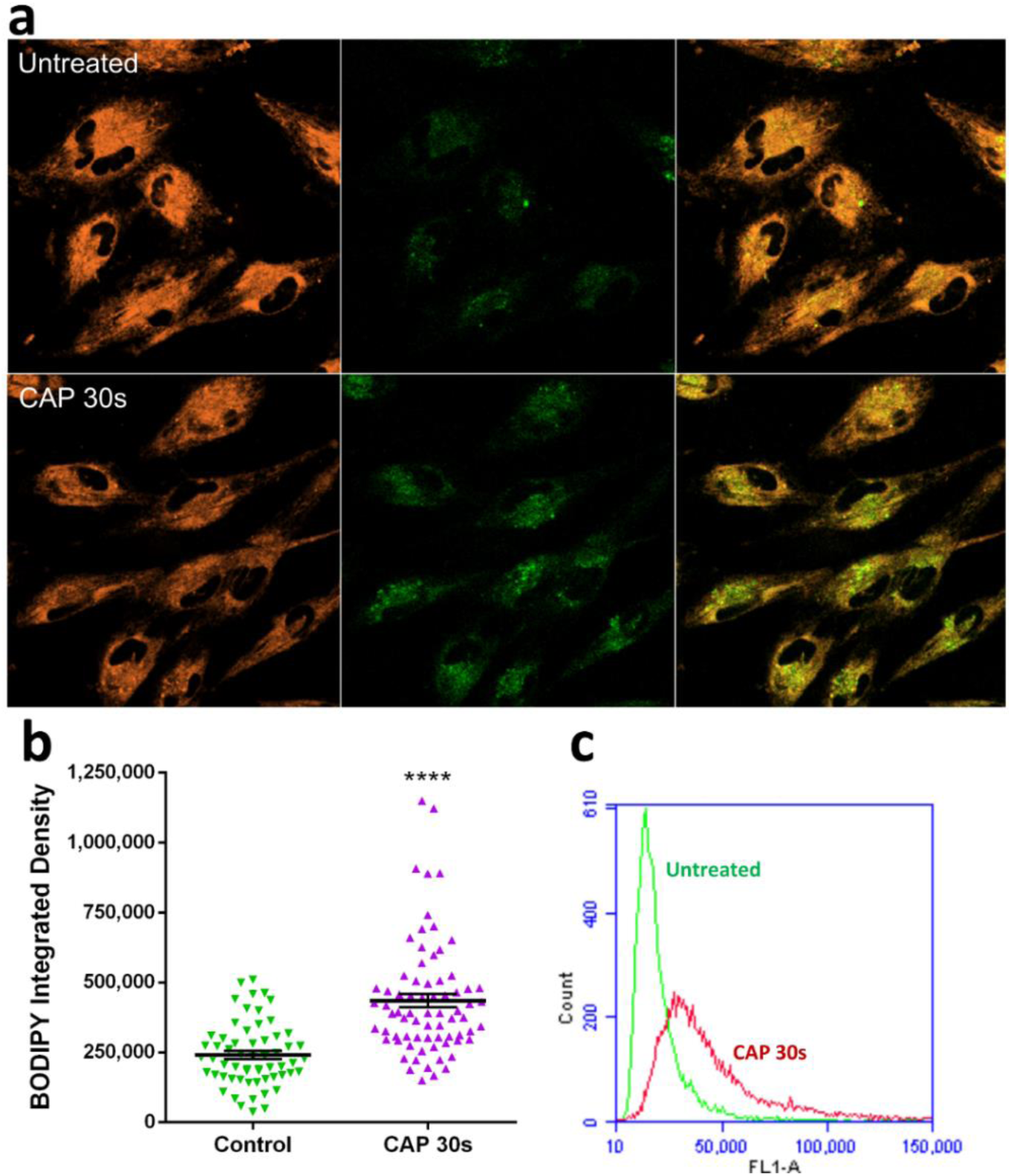
C11-BODIPY (581/591) staining shows lipid peroxidation and membrane trafficking inside the cell to lysosomes. (**a**) Confocal imaging of C11-BODIPY loaded cells; Orange, reduced form of C11-BODIPY; Green, oxidised form of C11-BODIPY. (**b**) The fluorescent integrated density of oxidised BODIPY was quantified using ImageJ. The statistical significance was assessed by one-way ANOVA with Tukey’s multiple comparison post-test (*P<0.05, **P<0.01, ***P<0.001, ****p<0.0001), n≥50, see Supplementary Table S1 for original data. (**c**) Fluorescence level of intracellular oxidised BODIPY was measured via Flow cytometry, left curve (green, untreated cells), right curve (red, CAP-treated cells).

### Tracking AuNPs in endosomes and lysosomes and effects of CAP treatment on endocytosis of AuNPs

A membrane repair mechanism has been described for cells^44, 45^. Through rapid endocytosis, cells can quickly remove damaged regions of membranes from the cell surface. These impaired membranes can be trafficked into endosomes and finally to lysosomes. In our case, the rapid endocytosis may contribute to the increased uptake of AuNPs or other compounds into cells following CAP treatment.

To test this hypothesis, CellLight™ Early Endosomes-RFP, BacMam 2.0 and CellLight™ Late Endosomes-RFP, BacMam 2.0 were used to further determine whether the route of uptake of AuNPs after CAP treatment was associated with endocytosis. Rab5a and Rab7a chimeras tagged with RFP were transfected and expressed inside cells. After overnight incubation, Rab5a-RFP and Rab7a-RFP were used to specifically track early endosomes and late endosomes, respectively^46^. In Supplementary Figure S6, the white arrows identify examples of co-localisation of AuNPs (red) with either early (left) or late (right) endosomes (orange). Lysosomes (green) were also counterstained. We have previously demonstrated that AuNPs accumulate in lysosomes 24 hours after 30 s, 75 kV CAP treatment using confocal imaging and 3D-image construction^11^. We demonstrate here that that AuNPs enter U373MG cells mainly through endocytosis, which can be identified first in early and late endosomes after CAP treatment and eventually accumulate in lysosomes (Supplementary Figure S6).

To monitor the immediate rate of change of endocytosis after CAP treatment, U373MG cells were treated with CAP for 30 s, 75 kV, then incubated with transferrin conjugated with Alexa Fluor™ 546, which is used as an early endosome marker, for 5 min, then fixed with 4% PFA. Transferrin specifically binds to the Transferrin receptor on the cell membrane to deliver Fe^3+^ atoms via receptor-mediated endocytosis. Iron-carrying transferrin then releases iron in the acidic environment of lysosome and will be recycled to the cell membrane. Therefore, early endosomes, including recycling endosomal pathways, were marked with red fluorescence within the confocal image (Figure 4a). As seen from the imaging, within 5 mins after CAP treatment, the number of transferrin-containing endosomes was greater compared to the control group. To statistically analyse the increase of endosomes induced by CAP treatment, more than 50 cells in each group were analysed using the ImageJ software. Quantification of transferrin uptake confirmed a significant increase of endosomes 5 min after CAP treatment (Figure 4b, p<0.0001).

**Figure 4.**
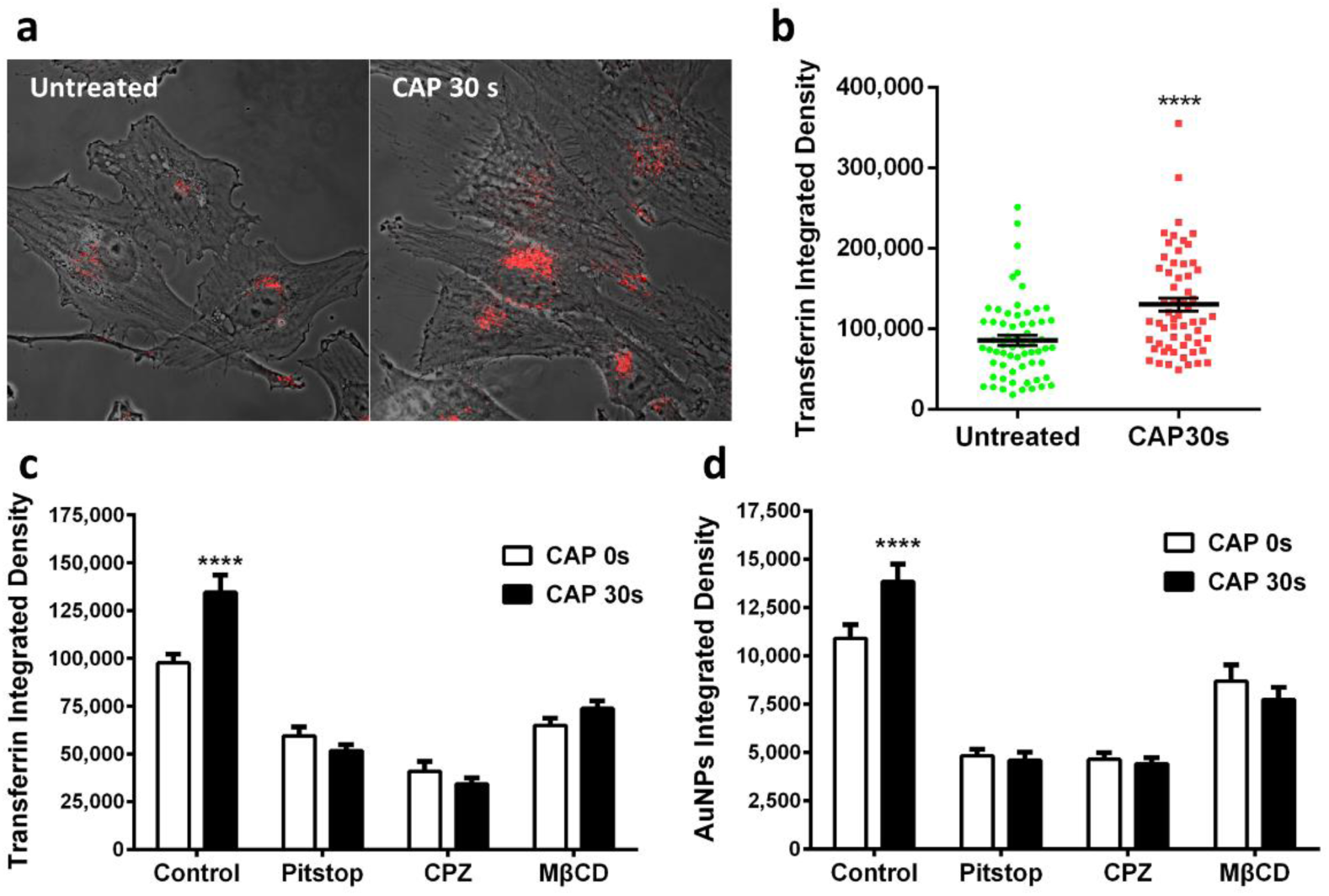
The CAP-induced endocytosis is clathrin-dependent. (**a**) After CAP treatment (0, 30 s), cells were loaded with transferrin-conjugated with Alexa Fluor™ 546 (red) for 5 min and fixed before observing with a confocal microscope. (**b**) The fluorescence level of transferrin was quantified using ImageJ, see Supplementary Table S2 for original data. (**c, d**) After incubation with various inhibitors as indicated, U373MG cells were treated with CAP for 0, 30 s at 75 kV and then loaded with transferrin for 5 min or 100 μg/ml AuNPs for 3 h respectively before observing using confocal microscopy, with the fluorescence integrated densities quantified using ImageJ. The statistical significance in (**b, c, d**) was assessed by one-way ANOVA with Tukey’s multiple comparison post-test (*P<0.05, **P<0.01, ***P<0.001, ****p<0.0001), n≥50.

Endocytosis is typically subdivided into four types, including clathrin-mediated endocytosis (CME), caveolae-mediated endocytosis, macropinocytosis and phagocytosis. As seen in Table 2, Pitstop, chlorpromazine (CPZ), Methyl-β-cyclodextrin (MβCD), filipin, genistein and amiloride are specific inhibitors suppress certain types of endocytosis and were used in this study to delineate the specific endocytic pathway activated by CAP.

**Table 2.**
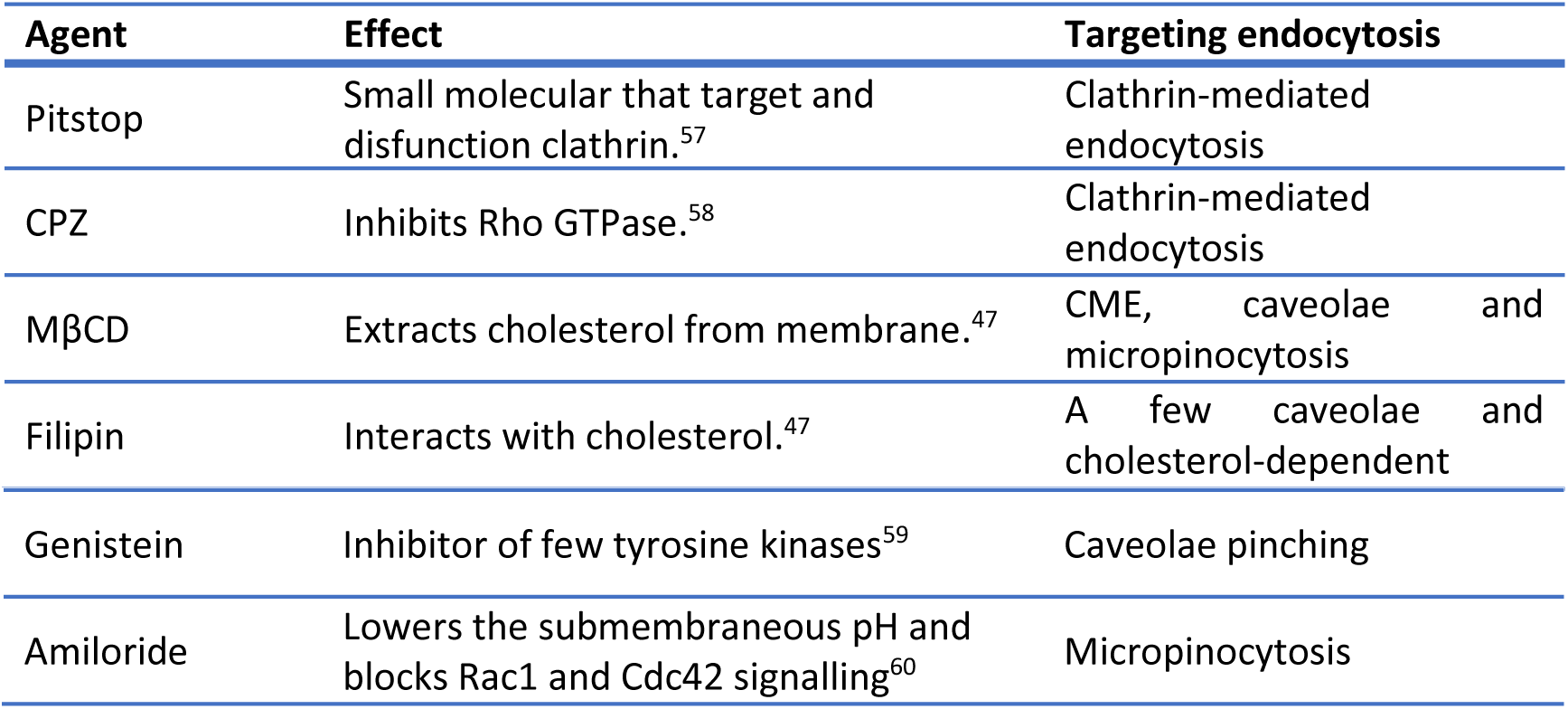
Inhibitors used to inhibit endocytosis in this research. ^47^.

| Agent | Effect | Targeting endocytosis |
| --- | --- | --- |
| Pitstop | Small molecular that target and disfunction clathrin.<sup>57</sup> | Clathrin-mediated endocytosis |
| CPZ | Inhibits Rho GTPase.<sup>58</sup> | Clathrin-mediated endocytosis |
| MβCD | Extracts cholesterol from membrane.<sup>47</sup> | CME, caveolae and micropinocytosis |
| Filipin | Interacts with cholesterol.<sup>47</sup> | A few caveolae and cholesterol-dependent |
| Genistein | Inhibitor of few tyrosine kinases<sup>59</sup> | Caveolae pinching |
| Amiloride | Lowers the submembraneous pH and blocks Rac1 and Cdc42 signalling<sup>60</sup> | Micropinocytosis |

U373MG cells were pre-incubated with inhibitors as indicated in the Methods section, then treated with CAP at 75 kV for 30 s. After treatment, cells were then incubated with fresh culture medium containing 100 μg/ml AuNPs for 24 hours or stained with transferrin-conjugated with Alexa Fluor™ 546 for 5 minutes then fixed before observing under a confocal microscope. To further quantify the fluorescence intensities of AuNPs and transferrin, more than 50 cells for each group were analysed using the ImageJ software (Supplementary Table S3 for original data). More transferrin-labelled endosomes (Figure 4c) and more AuNP (Figure 4d) were evident after CAP treatment of cells. Clathrin-inhibitors and MβCD-induced cholesterol depletion decreased the number of transferrin labelled endosomes and AuNP-uptake following CAP treatment (Figure 4c, d). Conversely, caveolae-specific inhibitors did not lead to any significant inhibition of Transferrin or AuNP endocytosis in cells after CAP treatment (Supplementary Figure S4). Macropinocytosis can simultaneously and un-specifically uptake all substances, including AuNPs and transferrin, in the fluid phase area of the cellular invagination of membrane^47^. As seen in Supplementary Figure S4, the accumulation of AuNPs and fluorescence level of transferrin were simultaneously decreased in amiloride-inhibited groups with or without CAP treatment and there remained a significant difference between CAP-treated and untreated groups.

To further confirm that clathrin-mediated endocytosis played the main role in CAP-accelerated cellular uptake, MISSION® Endoribonuclease-prepared siRNA (esiRNA) against human Clathrin heavy chain 1 (CLTC) was used to disrupting endocytosis mediated by clathrin coated pit formation. Cells were preincubated with MISSION® esiRNA (human CLTC) and MISSION® siRNA transfection reagent for 24 h, then treated with CAP for 0-30 s at 75 kV, as indicated in Figure 5, and then incubated with 100 μg/ml AuNPs in medium for 3 h and observed by confocal microscopy. As seen in Figure 5a-d, a significant decrease in AuNPs was observed in clathrin-silenced cells compared to the control groups. Following CAP treatment, we did not observe any increase in AuNP uptake in clathrin-silenced cells. To further quantify the uptake of AuNPs affected by clathrin-silencing and CAP treatment, more than 60 cells in each group were analysed using ImageJ software (Supplementary Table S4 for original data). Figure 5e represents this data and demonstrates that clathrin silencing inhibited more than 50% of the baseline AuNP uptake; moreover, no increase of AuNP uptake was measured following CAP treatment in the clathrin-silenced group. Together, this confirms that clathrin-mediated endocytosis played an important role in AuNP uptake, and accelerated endocytosis following CAP treatment was clathrin dependent.

**Figure 5.**
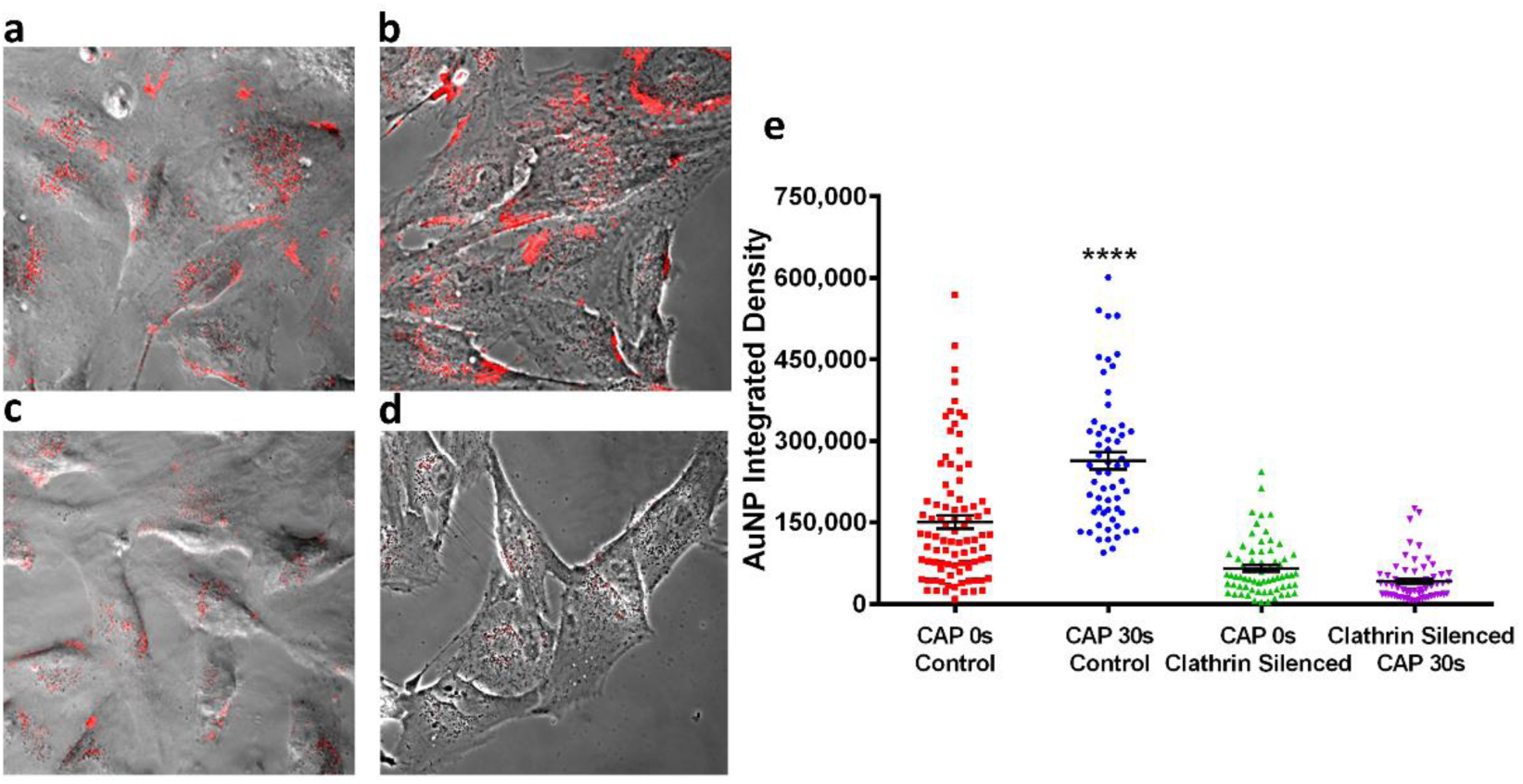
Clathrin silencing inhibits AuNPs uptake and CAP-induced endocytosis. (**a-d**) CAP-untreated cells without silencing, CAP-treated cells without silencing, CAP-untreated cells with silencing of clathrin heavy chain, CAP-treated cells with silencing of clathrin heavy chain. AuNPs are present as red fluorescence inside cells. (**e**) After incubation with esiRNA targeting expressing of clathrin heavy chain, U373MG cells were treated CAP for 0, 30 s at 75 kV and then loaded with 100 μg/ml AuNPs for 3 h before observing under confocal microscope, the fluorescence level was quantified using ImageJ. The statistical significance was assessed by one-way ANOVA with Tukey’s multiple comparison post-test (*P<0.05, **P<0.01, ***P<0.001, ****p<0.0001), n≥50.

## Discussion

The cytoplasmic membrane separates and protects the cellular interior from the exterior environment and provides specific and efficient exchange channels for the remaining intercellular balance and cell viability. Therefore, the integrity of the membrane is vital for all cells. Mammalian cells have developed efficient membrane repair mechanisms that can recover and reseal an injured cytoplasm membrane quickly to retain cell viability. Although investigations of the precise membrane repairing mechanisms have been limited, four possible mechanisms, including patch, tension reduction and more recently exocytosis/endocytosis and budding repair mechanisms were recently proposed^45^. The study of cytoplasmic membrane repair usually employs bacterial pore-forming toxins, such as Streptolysin O, to create mechanical injuries on membranes^45^. Meanwhile, lipid peroxidation is a complex process that damages cellular membrane structure and function, which is believed to link to numerous human diseases and aging, including Alzheimer diseases, dementia, Huntington, Parkinson, and traumatic injuries, under presence of oxidative stress^48, 49^. Many studies have shown that lipid peroxidation have various significant effects to cellular membranes, such as increase of membrane permeability ^50–52^, alteration of the lipid order and membrane fluidity^51, 53–55^ and activity change of membrane proteins^55–58^. However, there remains a paucity of literature identifying mechanisms of lipid peroxidation-related membrane damage repair, and the mechanisms of oxidised membrane repair remain unknown.

CAP is known to generate reactive species and thereby cause lipid peroxidation of cells. In this research, we explored CAP-induced lipid peroxidation using low dose DBD CAP treatment and studied the possible mechanisms of accelerated cellular uptake of AuNPs following CAP-induced oxidative membrane damage.

As seen in Figure 1, the uptake and accumulation of AuNPs into U373MG cells has been modelled for further investigation. To faithfully reproduce the experimentally observed results, a new independent uptake rate was added in the model (equation 5). This numerical model indicates that CAP treatment may introduce a new uptake route, which has now been determined as an independent membrane repairing, clathrin-mediated, endocytosis pathway.

As seen in Figure 2, we observed a relatively high level of reactive species generated in air during CAP treatment as detected by OES and O_3_ measurement. Several reactive species, such as ^•^OH, ONOO^−^ and O_3_, etc., generated in DBD CAP can induce lipid peroxidation in U373MG cells. Although there is no detectable emission of NO^•^, O, NO_x_ and low intensities of ^•^OH and N_2_ ^+^, which may due to interaction with relatively high quantities of O_3_, Figure 2a, indicates that there are high intensities of excited nitrogen molecules which can induce the generation of nitrated or peroxidised lipid. For example, it has been demonstrated that the reaction of excited N_2_ with H_2_ and H_2_O molecules can lead to the formation of H and ^•^OH products, respectively, which can induce lipid peroxidation^50^. Furthermore, as seen in Figure 2b, the significant increase of O_3_ levels during low dose cold plasma treatment shows the ability to peroxidise membrane lipids. The generated RONS in the CAP-treated culture medium, including hydrogen peroxide, nitrite and nitrate, were also quantitatively measured (As seen in Figure 2c). By comparison with our previous study^38, 42^, after the same low does CAP treatment (30 s, 75 kV), the levels of generated H_2_O_2_, NO^−^_2_, and NO^−^_3_ in culture medium are relatively low cytotoxic to U373MG cells, but still are able to induce lipid peroxidation.

From another perspective, the CAP treated group also showed significantly greater H_2_DCFDA fluorescence levels compared to the control group (Figure 2d), which confirmed the CAP-induced generation of reactive oxygen species inside cells. H_2_DCFDA is a cell permeant reagent which is fluorescent in its oxidised form, after interacting with hydroxyl, peroxyl and other reactive oxygen species. It has been widely reported that there are several by-products generated during lipid peroxidation, which can be used for specific markers, including malondialdehyde (MDA), 4-hydroxynonenal (HNE), 4-oxo-2-nonenal (ONE), and acrolein^51^.

The TBARS assay has been widely applied in food and biomedical research for detecting lipid peroxidation, as it can precisely and specifically measure the cellular level of MDA^52^. Figure 2e directly confirmed the CAP-induced lipid peroxidation, as a significant increase of MDA after CAP treatment appeared compared to the control group.

It has been shown that CAP treatment can alter membrane structures, which may be partly due to the reactive species-caused lipid peroxidation^42, 51, 53, 54^. However, the cell membrane remains PI impermeable after exposing to 75 kV CAP for 30 s, which demonstrates that the oxidised membrane may remain mechanically intact and the U373MG cells retain viability after a low dose of CAP treatment (30 s, 75 kV) (Figure 2f), which aligns with the results indicating low concentration of RONS in the CAP-treated medium (Figure 2c). As seen in Figure 3, there was a significant increase in the green fluorescence level of C11-BODIPY observed, which demonstrated that CAP treatment (30 s, 75 kV) induced significant peroxidation of BODIPY fatty acid, leading to a shift of the emission peak. There was also a significant increase of the vesicle structure marked by oxidised green C11-BODIPY tracked inside the U373MG cells, which may be the peroxidised membrane trafficked inside cells via endosomes (Figure 3a). The increase of endosomes was also confirmed using transferrin conjugated with Alexa Fluor™ 546. As seen in Figure 4a, b, the number of transferrin-marked endosomes within cells was significantly increased just 5 min after CAP treatment (30 s, 75 kV).

Therefore, we propose that low dose CAP treatment can cause non-lethal cytoplasmic membrane damage which triggers a rapid membrane repair system. The increased endocytosis induced by membrane repair then accelerates the uptake of AuNPs into U373MG cells.

To further elaborate on this hypothesis, the uptake of AuNPs into cells was tracked using CellLight™ Early Endosomes-RFP, BacMam 2.0, CellLight™ Late Endosomes-RFP, BacMam 2.0. and LysoTracker™ Green DND-26. In Supplementary Figure S6, after incubation for 3 hours, the white arrows show examples of AuNPs located in the early/late endosomes and lysosomes. Our previous work demonstrated that AuNPs accumulate in lysosomes, with significantly more accumulation in the case of CAP-treated cells, after 24 h incubation using confocal imaging and 3D-image construction^11^. Supplementary Figure S6 demonstrates the co-localisation of AuNPs with early and late endosome respectively, which shows a tendency of AuNPs uptake to enter the periphery region of the cells through endocytosis, then gather at the central zone of the cells via endosome trafficking into lysosomes. Therefore, it further confirms that U373MG cells mainly accumulate AuNPs into lysosomes via uptake into early then late endosomes after CAP treatment (30 s, 75 kV).

Various endocytosis inhibitors, AuNPs tracking and transferrin staining were used to determine the specific endocytosis pathways activated by CAP treatment. Transferrin is an essential iron-binding protein that facilitate iron-uptake in cells via transferrin receptor and Clathrin-mediated endocytosis. Figure 4c shows that the formation of transferrin-trafficking endosomes was inhibited in both groups with 0 and 30 s CAP treatment, after exposure to specific/non-specific clathrin inhibitors, including pitstop, CPZ and MβCD. CAP treatment no longer enhanced endocytosis when clathrin was inhibited. Tracking the accumulation of AuNPs after 24 hours incubation demonstrates the long-term effects retained post CAP treatment and various inhibitors (Figure 4d), which suggests similarly that CAP treatment can specifically activate clathrin-mediated endocytosis to enhance cellular uptake of AuNPs. Clathrin silencing combining CAP treatment further confirmed that the CAP-triggered endocytosis for membrane repairing is clathrin dependent (Figure 5).

In summary, we report that the enhanced uptake of AuNPs induced by CAP can be as a result of ROS-caused lipid peroxidation, leading to rapid cytoplasma membrane repairing via clathrin-dependent endocytosis. This contributes to our understanding of the cellular effects induced by CAP, especially membrane damage and endocytosis activation, which can be employed for efficient uptake of nanomaterials and pharmaceuticals into cells when combining CAP with cancer therapies. This mechanism of RONS-induced endocytosis will also be of relevance to researchers optimizing other cancer therapies that induce an increase in extracellular RONS.

## Methods

### Cell Culture and Gold Nanoparticle Treatment

U373MG-CD14, human brain glioblastoma cancer cells (Obtained from Dr Michael Carty, Trinity College Dublin) were cultured in DMEM-high glucose medium (Merck) supplemented with 10% FBS (Merck) and maintained in a 37 °C incubator within a humidified 5% (v/v) CO_2_ atmosphere. Gold nanoparticles were synthesised by trisodium citrate reduction of auric acid. 20nm sphere citrate-capped AuNPs were used to treat cells whose properties were determined in a previous study^11^. The gold colloid was concentrated to 2500 μg/ml then diluted in culture medium to 100 μg/ml which is non-toxic to U373MG cells. The culture medium containing 100 μg/ml AuNPs was then used to treat cells as indicated in the relevant figures.

### CAP Configuration and Treatment

The current research uses an experimental atmospheric dielectric barrier discharge (DBD) plasma reactor, which has been described and characterised in detail^33, 55^. All U373MG cells were treated within containers, which were placed in between two electrodes, at a voltage level of 75 kV for 30 s. Prior to CAP treatment, the culture medium was removed, and fresh culture medium was added into the cell culture container at 5% of the well working volume to prevent drying during treatment. Afterwards, fresh culture medium containing 100 μg/ml AuNPs or inhibitors as indicated was added to reach the well final working volume and incubated with cells at 37 °C for the indicated time.

### H_2_DCFDA Assay and Optical Emission Spectroscopy (OES) and Ozone measurement

H_2_DCFDA (Thermo Fisher Scientific) was used to detect ROS induced by CAP treatment. U373MG cells were seeded into the TC dish 35 standard (35×10mm, Sarstedt) at a density of 2×10^5^ cells/ml and incubated overnight to allow adherence. After washing twice with PBS, cells were incubated with 25 μM H_2_DCFDA in serum-free medium for 30 min at 37 °C. Cells were then washed with PBS twice, culture medium once and then treated with CAP at 75 kV for 30s. The fluorescence of H_2_DCFDA was then measured using flow cytometry.

Optical emission spectroscopy was carried out using an Edmund Optics CCD spectrometer with a spectral resolution of between 0.6 nm to 1.8 nm. The spectra were measured using BWSpec^TM^ software with a spectral range between 200 and 850 nm and were acquired every 7.5 s with an integration time of 1500 ms. Total relative intensity of each emission line was calculated using the integral of the area under each peak. EEDF was calculated using a line ratio method (N_2_ at 337 nm and N_2_ ^+^ at 391 nm)^56^. O_3_ was sampled using a standard Gastec sampling pump in conjunction with a Gastec detection tubes immediately after plasma discharge had ceased.

### Measurement of hydrogen peroxide, nitrite and nitrate concentrations

The concentrations of hydrogen peroxide, nitrite and nitrate were quantitatively measured in CAP-treated culture medium without phenol red. Concentrations of hydrogen peroxide, nitrite and nitrate were determined employing the TiSO_4_ assay, Griess reagent and 2,6-dimethylphenol (DMP) assay, respectively. The quantitative methods have been described in detail elsewhere^39^. For the treatment of culture medium, 150 μl of non-phenol red culture medium with or without 100 μg/ml AuNPs were treated with CAP at 75kV for 30 s in 35 mm dishes containing 80% confluent cells, then collected for measurement. To eliminate the effect of the culture medium on photometrical measurements, the results of CAP-treated groups were standardised with untreated culture medium.

### Lipid peroxidation

Thiobarbituric acid reactive substances (TBARS) assay was used to measure the lipid peroxidation induced by CAP in U373MG cells. MDA was measured and normalized to total protein. Thiobarbituric acid (TBA), trichloroacetic acid, MDA were purchased from Merck. U373MG cells were seeded into TC Dish 150 (Standard, 150×20 mm, Sarstedt) at a density of 1×10^5^ cells/ml and incubated until confluence. Then the cells were treated with CAP for 30 s at 75 kV. After treatment, cells were further incubated for 24 h, and collected by trypsinisation, centrifuged (100 g for 5 min), and homogenized by sonication. 100 μl homogenate was mixed with 200 μl ice cold 10% trichloroacetic acid and incubated on ice for 15 min to precipitate protein, then centrifuged (2200 g for 15 min at 4 °C). 200 μl of each supernatant was then mixed with 200 μl 0.67% (w/v) TBA and incubate at 100 °C for 10 min. The samples were measured at 532 nm for MDA after cooling. The level of MDA was compared to a standard curve of 0-100 μM MDA, and normalized to total protein, which was measured in cell homogenates using the Bradford assay.

Lipid peroxidation sensor, C11-BODIPY (581/591) (Thermo Fisher Scientific) was used for *in-situ* detection and localization of the lipid peroxidation induced by CAP treatment. Oxidation of this lipophilic fluorophore results in a change of the fluorescence emission peak from ∼590 nm to ∼510 nm. 1mM stock solution was prepared in DMSO and cells were incubated in fresh culture medium containing 5 μM of the probe at 37 °C for 30 min in advance. Then the cells were washed with PBS twice and culture medium once. Afterwards, cells were treated with CAP for 30 s at 75 kV, incubated for 30 min, and observed using a Zeiss LSM 510 confocal laser scanning microscope as described later. The following excitation and emission settings were used: Excitation wavelength 1: 488 nm, emission filter 1: 500-560; Excitation wavelength 2: 568 nm, emission filter 2: 560-620.

### Flow cytometry

U373MG cells were seeded into a TC dish 35 standard (35×10 mm, Sarstedt) at a density of 2×10^5^ cells/ml and incubated overnight to allow a proper adherence. Cells were then loaded with C11-BODIPY and washed as described above (Lipid peroxidation). After treating with CAP, cells were incubated with fresh medium for 0.5h at 37 °C before collecting.

BD Accuri™ C6 Plus flow cytometry (BD Bioscience) was used to detect and track fluorescence of the lipid peroxidation sensor. To prepare aliquots, all floating and attaching cells were collected by trypsinisation and then washed twice with PBS. All liquids, including medium, PBS and trypsin-cell suspension, were collected into one tube, then centrifuged at 100g for 5 min. 2 ml of fresh PBS was used to resuspend the cell tablet for assessment by flow cytometry. For the measurement, a 488 nm laser was used for excitation, and 10,000 gated events were collected. Green fluorescence (oxidised dye) and red fluorescence (non-oxidised dye) was measured using an FL1 standard filter (533/30 nm) and FL2 standard filter (585/40 nm), respectively.

For Propidium iodide (PI) staining, U373MG cells were seeded into TC Dish 35 at a density of 2×10^5^ cells/ml and incubated overnight to allow adherence. Cells were then exposed to CAP 75kV for 30s. Afterwards, cells were incubated at 37 °C for 30 minutes, and collected by trypsinisation, resuspended into 1ml PBS. Resuspended cells were stained with 1µg/ml PI for 5 minutes. The fluorescence of PI was then measured using BD Accuri™ C6 Plus flow cytometry at FL2 (585/40nm) standard filter.

### Inhibitor Studies

To inhibit various endocytic pathways, cells were pre-incubated with Pitstop (12.5 μM, 5 min) chlorpromazine (10 μg/ml, 10 min), filipin (5 μg/ml, 30 min), genistein (200 μM, 30 min), amiloride (50 μM, 30 min) and methyl-β-cyclodextrin (10 mM, 30 min) in culture medium for the time indicated, at 37 °C. After inhibiting treatment, the culture medium was removed during CAP treatment (75 kV, 30 s), prewarmed fresh culture medium containing 100 μg/ml AuNPs was then added immediately to the dishes and incubated for 3 h before observing using a Zeiss LSM 510 confocal laser scanning microscope.

Transferrin conjugated with Alexa Fluor™ 546 was used to determine the change of early endosomes induced by CAP combining various endocytosis inhibitors. After the inhibiting and CAP treatments indicated above, the cells were incubated in prewarmed fresh medium for 3 h, then incubated with 25 μg/ml transferrin in medium for 5 min. Afterwards, cells were fixed with 4% PFA and then observed using confocal microscopy. The details of the confocal microscope are described in following section.

### Clathrin Silencing

MISSION® esiRNA (human CLTC) and MISSION® siRNA transfection reagents were purchase from Merck. 50,000 U373MG cells were seeded into each 35 mm glass-bottom dishes (Greiner Bio-One) and incubated overnight. 1.2 ul esiRNA stock was mixed with 20 ul of transfection reagent in 400 μl serum-free medium and incubated for 15 minutes at room temperature. In glass-bottom dishes, previous medium was replaced with 1 ml of prewarmed fresh medium with serum. The siRNA/transfection reagent solution was then added onto the cells and homogenized to final volume of 1.2 ml. Afterwards, U373MG cells were incubated at 37 °C for 24 h and then incubated with fresh medium containing 100 μg/ml AuNPs for 3 h before observing using confocal microscopy.

### Endocytosis Tracking and Cell Imaging

Early endosomes, late endosomes, lysosomes were demonstrated using the CellLight™ Early Endosomes-RFP, BacMam 2.0, the CellLight™ Late Endosomes-RFP, BacMam 2.0 and the LysoTracker™ Green DND-26, respectively (Thermo Fisher Scientific). 35 mm glass-bottom dishes (Greiner Bio-One) were used as containers for confocal imaging. Cells were seeded at a density of 1×10^5^ cells/ml and incubated for 16 h. For early and late endosome marker, 2 μl of BacMan 2.0 reagent per 10,000 cells was added in fresh medium and incubated with cells at 37 °C for 16 h. Then the cells were treated with CAP for 30s and incubated with fresh medium containing 0-100 μg/ml AuNPs for 3 or 24 h at 37 °C as indicated. Afterward, the samples were observed using a Zeiss LSM 510 confocal laser scanning microscope. For lysosome tracking, cells were seeded for 16 h, treated with CAP for 30s, then incubated with AuNPs for 3 or 24 h. Afterward, cells were washed twice with PBS and incubated with medium containing 50 nM LysoTracker for 5 min at 37 °C. Cells were then washed once with PBS loaded with phenol red (Merck) and observed using a Zeiss LSM 510 confocal laser scanning microscope. The corresponding filter settings were as follows. AuNPs, excitation wavelength: 633 nm, reflection filter: 649-799 nm; Transferrin conjugated with Alexa Fluor™ 546, excitation wavelength: 568 nm, emission filter: 580-630 nm; CellLight™ Early Endosomes-RFP, BacMam 2.0, excitation wavelength: 568 nm, emission filter: 580-630 nm; CellLight™ Late Endosomes-RFP, BacMam 2.0, excitation wavelength: 568 nm, emission filter: 580-630 nm; LysoTracker™ Green DND-26, excitation wavelength: 488 nm, emission filter: 505-530 nm. Plan-Apochromat 63x/1.4 Oil Ph3 was used as objective for all samples. To analyse the fluorescent intensity, around 50 cells were randomly imaged for each treatment condition, then those images was analysed using the ImageJ and Prism 6 (GraphPad Software).

### Statistical Analysis

At least triplicate independent tests were carried out for each data point, unless indicated otherwise. Error bars of all figures are presented using the standard error of the mean (S.E.M). Prism 6 (GraphPad Software) was used to carry out curve fitting and statistical analysis. Two-tailed P values were used and the Alpha for all experiments is 0.05. The significance between data points was verified using one-way ANOVA and two-way ANOVA with Tukey’s multiple comparison post-test, as indicated in figures (*P<0.05, **P<0.01, ***P<0.001, ****p<0.0001).

## Acknowledgements

This work is supported by TU DUBLIN Fiosraigh Research Scholarship programme (Z.H., E.M., K.L., L.S., S.G., M.M.), Science Foundation Ireland Grant Numbers 14/IA/2626 (P.B., H.B., P.C. and J.C.) and 16/BBSRC/3391.

## Author Contributions

Z.H., F.T., P.C. and J.C. conceived the project and designed the experiments. Z.H., H.B., E.M., K.L., L.S., S.G., S.W.N. and M.M. performed the experiments, collected and analysed the data. Z.H., F.T., B.T., P.C, P.B, H.B. and J.C. co-wrote the paper. All authors discussed the results and reviewed the manuscript.

## Supplementary Materials

**Supplementary Figure S1.**
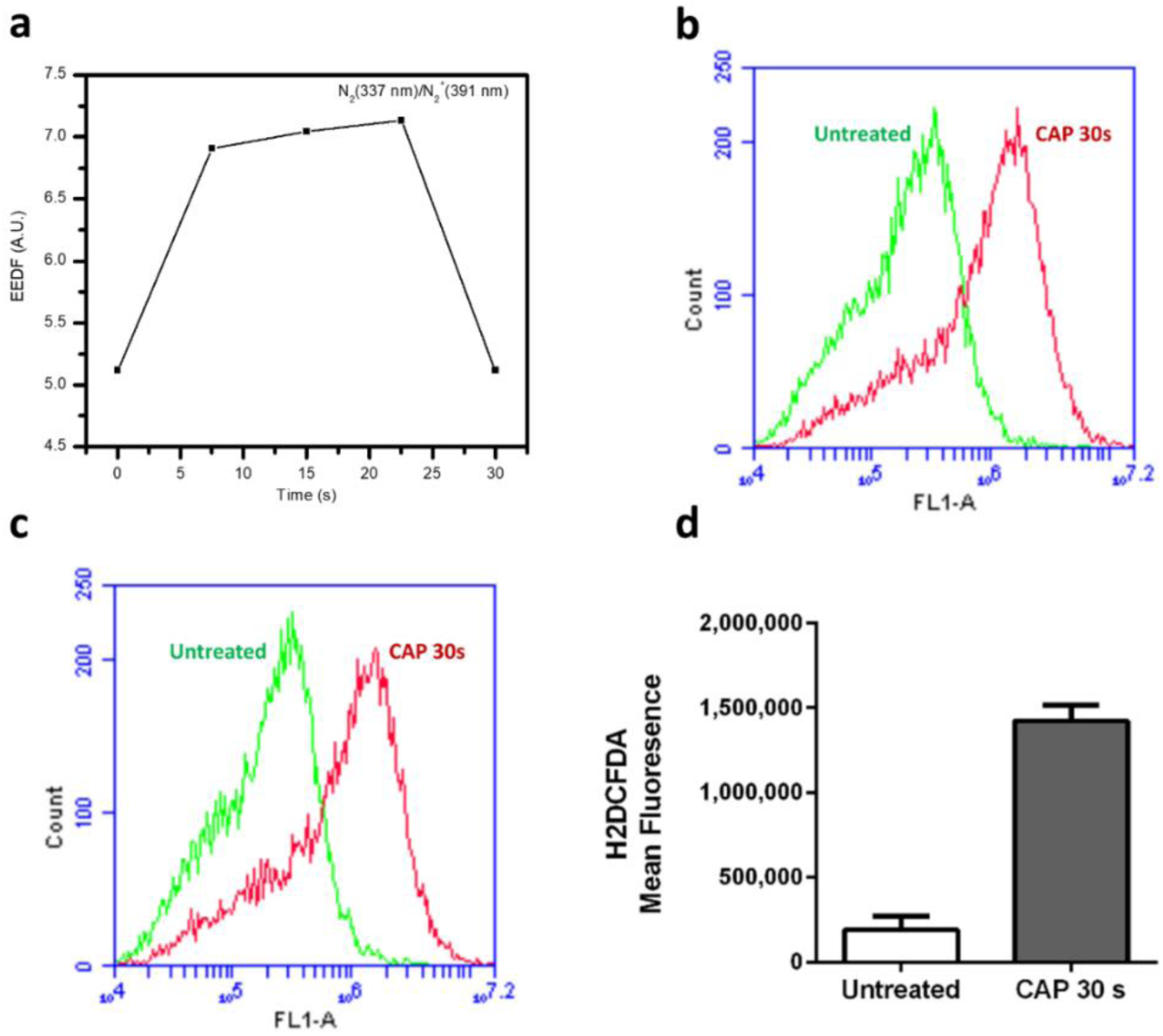
(a) Electro energy distribution function (EEDF) of CAP; Fluorescence level of intracellular oxidised H_2_DCFDA was measured via Flow cytometry, two replicas (b, c) and the analysed average of mean FL1-A value (d).

**Supplementary Figure S2.**
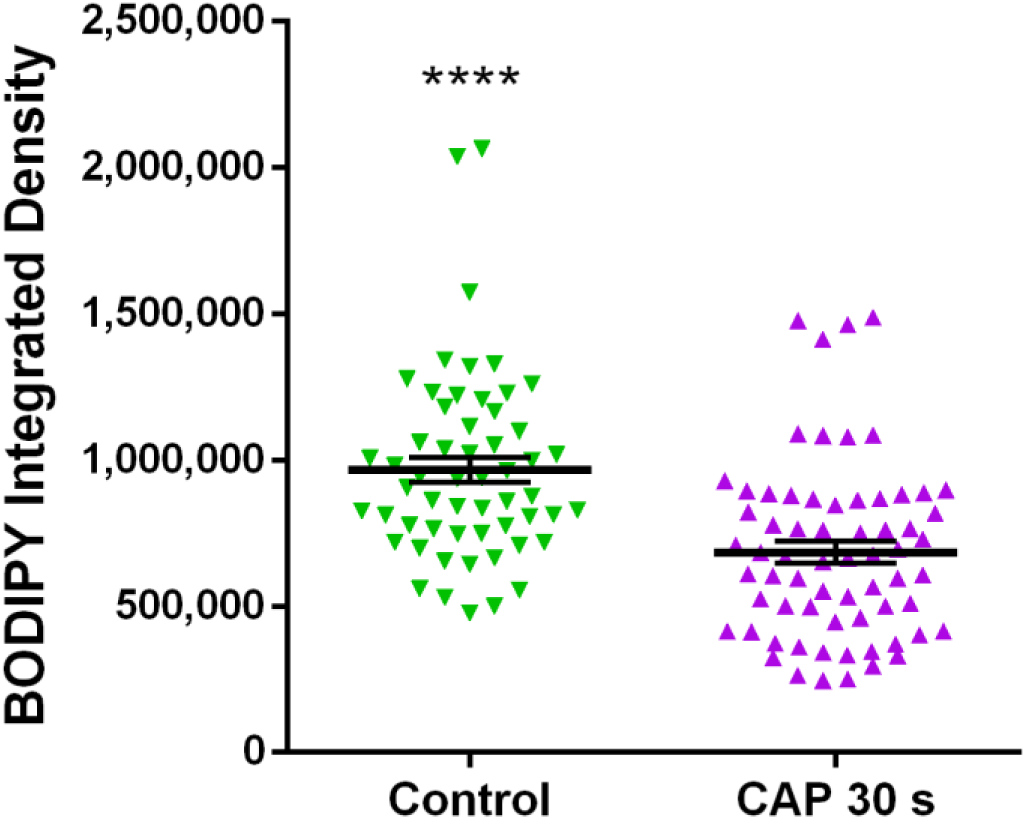
The fluorescent integrated density of non-oxidised BODIPY was quantified using ImageJ. The statistical significance was assessed by one-way ANOVA with Tukey’s multiple comparison post-test (*P<0.05, **P<0.01, ***P<0.001, ****p<0.0001), n≥50, see Supplementary Table S1 for original data

**Supplementary Figure S3.**
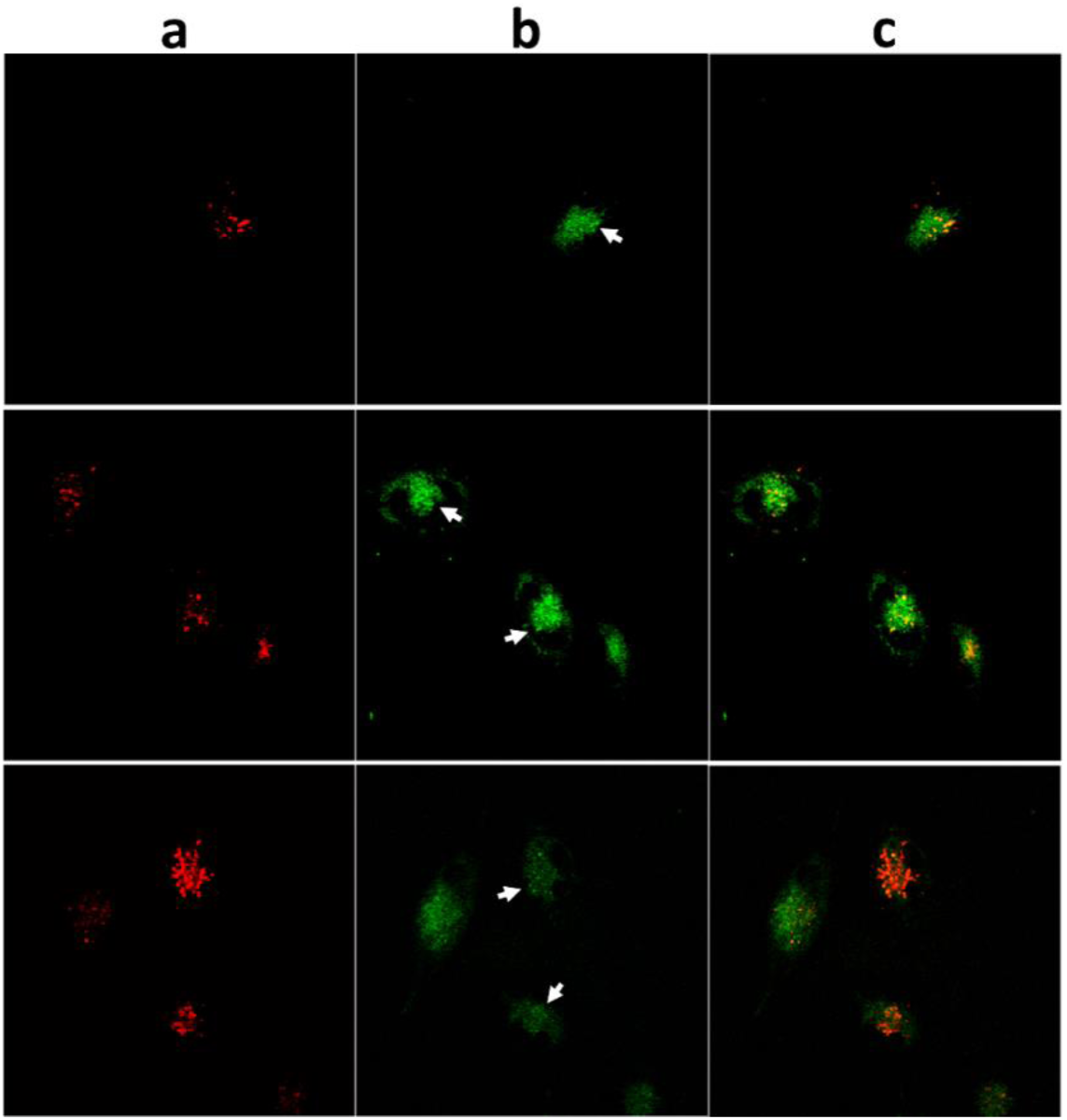
The U373MG cells were loaded with C11-BODIPY (green), then co-stained with and LysoTracker™ Deep Red (red) after CAP treatment (30 s, 75 kV) and observed under confocal microscope, three images are presented in the figure. White arrows point out examples of co-localisation between oxidised C11-BODIPY and lysosomes.

**Supplementary Figure S4.**
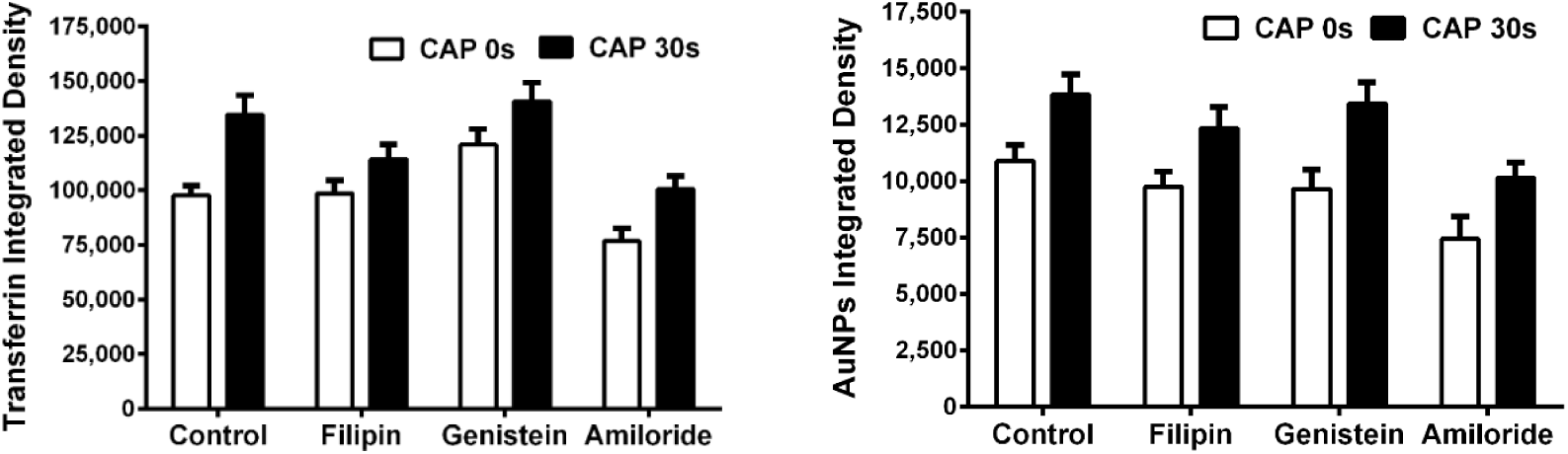
After incubation with various inhibitors as indicated, U373MG cells were treated CAP for 0, 30 s at 75 kV and then loaded with transferrin for 5 min or 100 μg/ml AuNPs for 3 h respectively before observing under confocal microscope, then the fluorescence integrated densities were quantified using ImageJ. The statistical significance was assessed by one-way ANOVA with Tukey’s multiple comparison post-test (*P<0.05, **P<0.01, ***P<0.001, ****p<0.0001), n≥50.

**Supplementary Figure S5.**
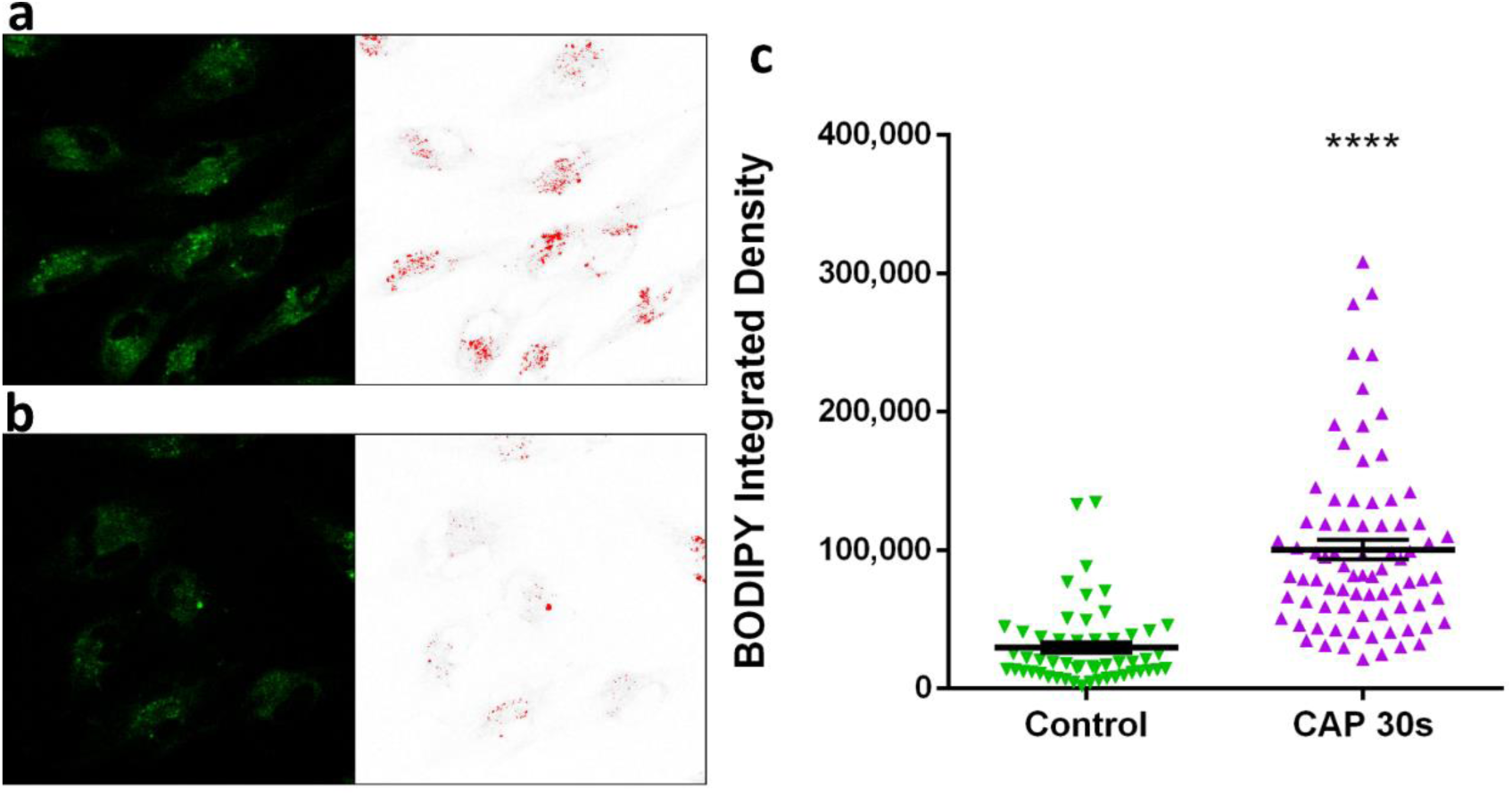
Oxidized C11-BODIPY emission was measured with same threshold to highlight vesicle-like condensed area. (a) Sample of CAP treated cells; (b) Sample of untreated cells; (c) With same threshold, the fluorescence integrated densities were quantified using ImageJ. The statistical significance was assessed by unpaired T-test. (*P<0.05, **P<0.01, ***P<0.001, ****p<0.0001), n≥50.

## Supplementary Discussion

The intensity of O_3_ production may explain why there was no detectable emission of NO, O, NO_x_ and low intensities of ^•^OH and N_2_^+^ in Figure 2a, as the free electrons were likely quenched before reaching higher energetic states by the interaction and formation of O_3_. Reaction mechanisms (1-4) can further explain how the formation of NO, NO_x_, and O was hindered. In these reactions, M is a third body atom or molecule that, in this case, may be N_2_* or O_2_.

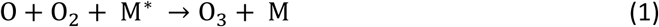

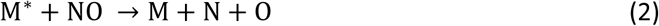

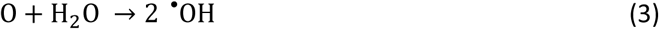

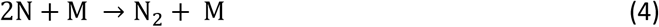

**Supplementary Figure S6.**
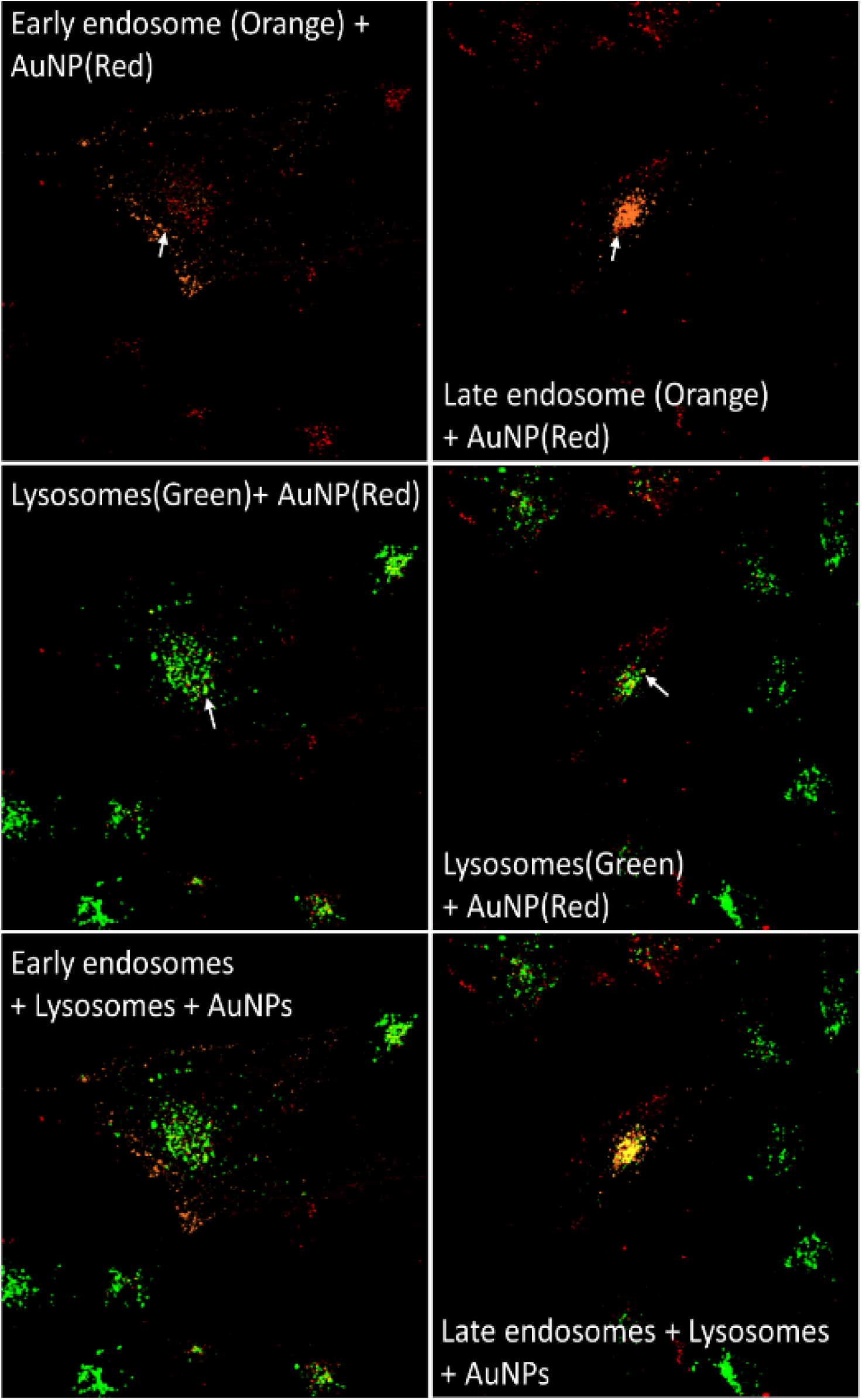
Uptake and subcellular localization of AuNPs observed by confocal microscopy. After CAP treatment (0, 30 s), U373MG cells were incubated with 100 μg/ml AuNPs for 3 h. Early, late endosomes and lysosomes were stained using CellLight™ Early/Late Endosomes-RFP (Marked as orange channel in images) and LysoTracker™ Green DND-26, respectively. The far-red emission of AuNPs (Red) was excited using 633 nm laser.

**Supplementary Table S1.**

| Integrated Density of BODIPY |  |  |  |
| --- | --- | --- | --- |
| Control. | CAP 30s. | Control. Oxidized (Figure 3b) | CAP 30s. Oxidized |
| 825441 | 764257 | 232430 | 417629 |
| 998949 | 760580 | 310618 | 342874 |
| 1060462 | 665879 | 327636 | 495228 |
| 859260 | 1089862 | 179182 | 477016 |
| 530275 | 765581 | 307326 | 426428 |
| 829356 | 360915 | 176954 | 448261 |
| 665668 | 551518 | 499637 | 293395 |
| 1278050 | 264406 | 510195 | 325466 |
| 2038367 | 596517 | 356545 | 273752 |
| 936308 | 760052 | 292490 | 464481 |
| 1259645 | 332197 | 407417 | 167146 |
| 1573896 | 866259 | 281113 | 394278 |
| 1019896 | 499450 | 227051 | 302926 |
| 963641 | 296252 | 181733 | 221856 |
| 862036 | 846493 | 189916 | 454239 |
| 812204 | 881837 | 248568 | 374735 |
| 1113481 | 861837 | 300329 | 373363 |
| 748271 | 778087 | 273441 | 505987 |
| 1165023 | 524382 | 211968 | 292939 |
| 764072 | 867958 | 282522 | 388750 |
| 1232175 | 508610 | 317893 | 597744 |
| 1007005 | 595270 | 161873 | 295272 |
| 982070 | 610424 | 155628 | 306654 |
| 1320705 | 445595 | 458594 | 390399 |
| 946150 | 1463426 | 399135 | 890896 |
| 1342500 | 708723 | 462811 | 741382 |
| 653886 | 888079 | 196388 | 524934 |
| 499775 | 373856 | 190853 | 306468 |
| 875556 | 1477152 | 153302 | 390161 |
| 1037540 | 653049 | 163432 | 651081 |
| 906223 | 1078557 | 142877 | 187065 |
| 1097569 | 675892 | 206789 | 906811 |
| 1207059 | 402444 | 37404 | 312273 |
| 718964 | 1083730 | 109317 | 617780 |
| 559771 | 930197 | 236604 | 468648 |
| 475472 | 898191 | 142135 | 296259 |
| 747381 | 331716 | 270552 | 389688 |
| 807312 | 532198 | 82436 | 195602 |
| 707968 | 817839 | 84652 | 506283 |
| 840742 | 414865 | 168778 | 451699 |
| 777642 | 749632 | 364207 | 303635 |
| 1022745 | 879071 | 100734 | 431292 |
| 1228769 | 1084756 | 114519 | 701205 |
| 2064986 | 412708 | 49002 | 444981 |
| 697882 | 884861 | 273032 | 223330 |
| 1220345 | 566726 | 58729 | 423576 |
| 775782 | 667011 | 169249 | 569403 |
| 1182171 | 895365 | 184585 | 660069 |
| 812884 | 497905 | 198073 | 227918 |
| 940440 | 603690 | 154684 | 478087 |
| 555395 | 684168 | 441372 | 254274 |
| 1327438 | 371186 | 274210 | 362961 |
| 718786 | 502118 | 245082 | 398478 |
| 642920 | 415450 | 228162 | 325251 |
| 1050097 | 1489285 | 437520 | 346672 |
| 834466 | 250668 | 350830 | 479690 |
|  | 247114 | 199513 | 1122291 |
|  | 460969 | 184087 | 191401 |
|  | 345379 | 260743 | 335414 |
|  | 325823 | 234688 | 346672 |
|  | 342280 |  | 303883 |
|  | 694899 |  | 306629 |
|  | 607436 |  | 1149608 |
|  | 821919 |  | 150280 |
|  | 1413574 |  | 456893 |
|  | 729336 |  | 283002 |
|  |  |  | 269830 |
|  |  |  | 234559 |
|  |  |  | 524097 |
|  |  |  | 372300 |
|  |  |  | 626520 |
|  |  |  | 889340 |
|  |  |  | 691489 |

**Supplementary Table S2.**

| Integrated Density of Transferrin |  |
| --- | --- |
| Untreated | CAP30s |
| 110807 | 57027 |
| 66999 | 112562 |
| 125539 | 86679 |
| 25907 | 97764 |
| 18663 | 210017 |
| 36431 | 151799 |
| 55003 | 82789 |
| 58136 | 180386 |
| 108838 | 219322 |
| 203364 | 120825 |
| 38237 | 109507 |
| 106023 | 181812 |
| 153065 | 232475 |
| 46973 | 103867 |
| 231021 | 165690 |
| 126260 | 189547 |
| 58261 | 145942 |
| 53516 | 216243 |
| 28616 | 175271 |
| 169818 | 79758 |
| 119425 | 197285 |
| 69592 | 173238 |
| 24623 | 207385 |
| 126382 | 137586 |
| 109035 | 103140 |
| 251395 | 55164 |
| 90739 | 287846 |
| 76501 | 49045 |
| 76803 | 75313 |
| 71061 | 76832 |
| 70167 | 89317 |
| 71700 | 116572 |
| 29972 | 81825 |
| 28606 | 205643 |
| 87132 | 134411 |
| 33269 | 101051 |
| 95125 | 355132 |
| 80060 | 57270 |
| 64769 | 73021 |
| 106737 | 108349 |
| 74147 | 88346 |
| 79054 | 56330 |
| 25151 | 115220 |
| 40603 | 109490 |
| 72073 | 124859 |
| 124198 | 218535 |
| 77852 | 134532 |
| 103302 | 71167 |
| 109488 | 71050 |
| 58064 | 60830 |
| 75705 | 58081 |
| 27911 | 63689 |
| 86282 | 181942 |
| 117192 | 170011 |
| 32705 | 89790 |
| 120661 | 71784 |
| 129909 | 163049 |
| 106761 | 107358 |
| 90118 | 132343 |
| 39569 |  |
| 164849 |  |
| 84330 |  |

**Supplementary Table S3.**
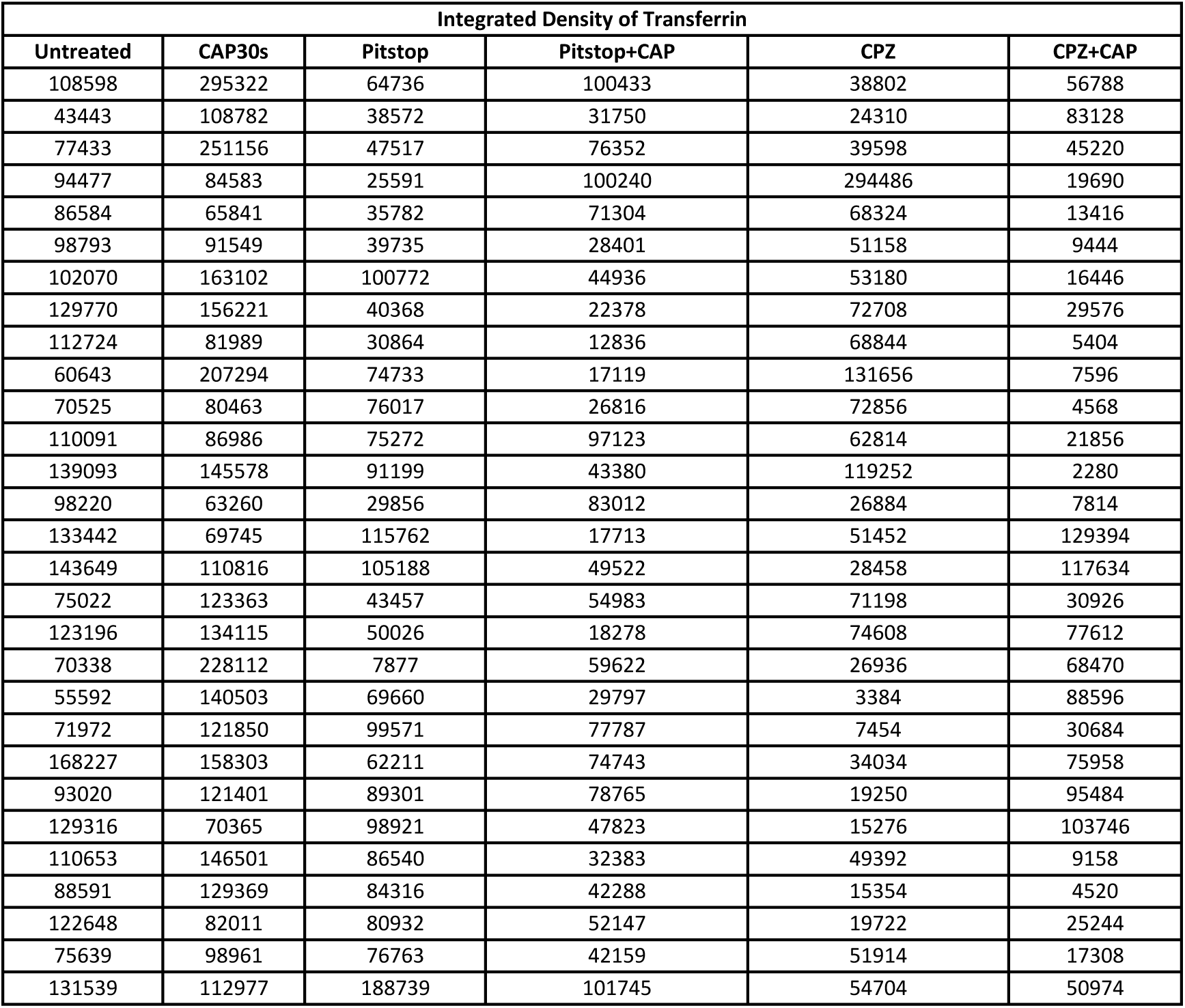
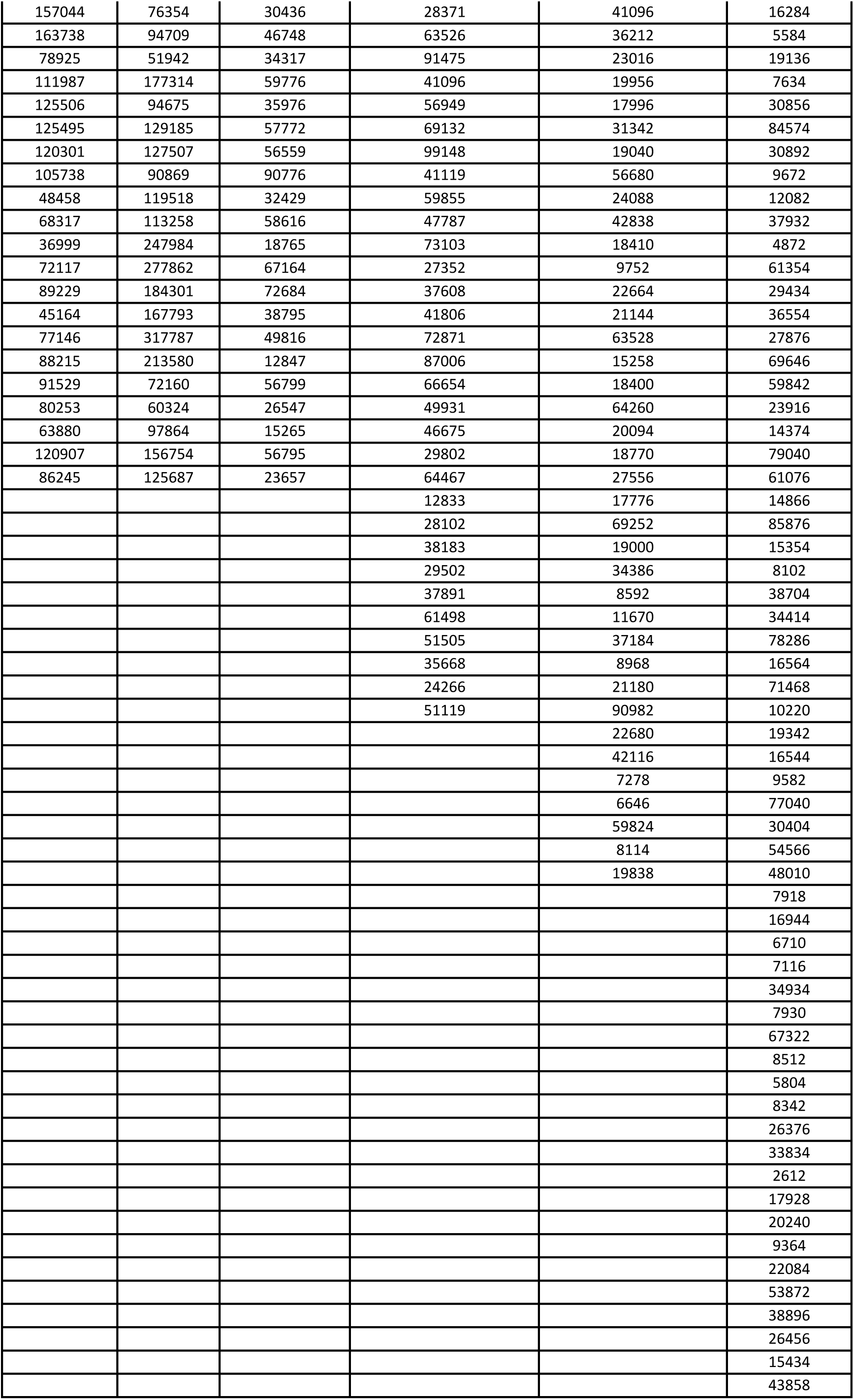
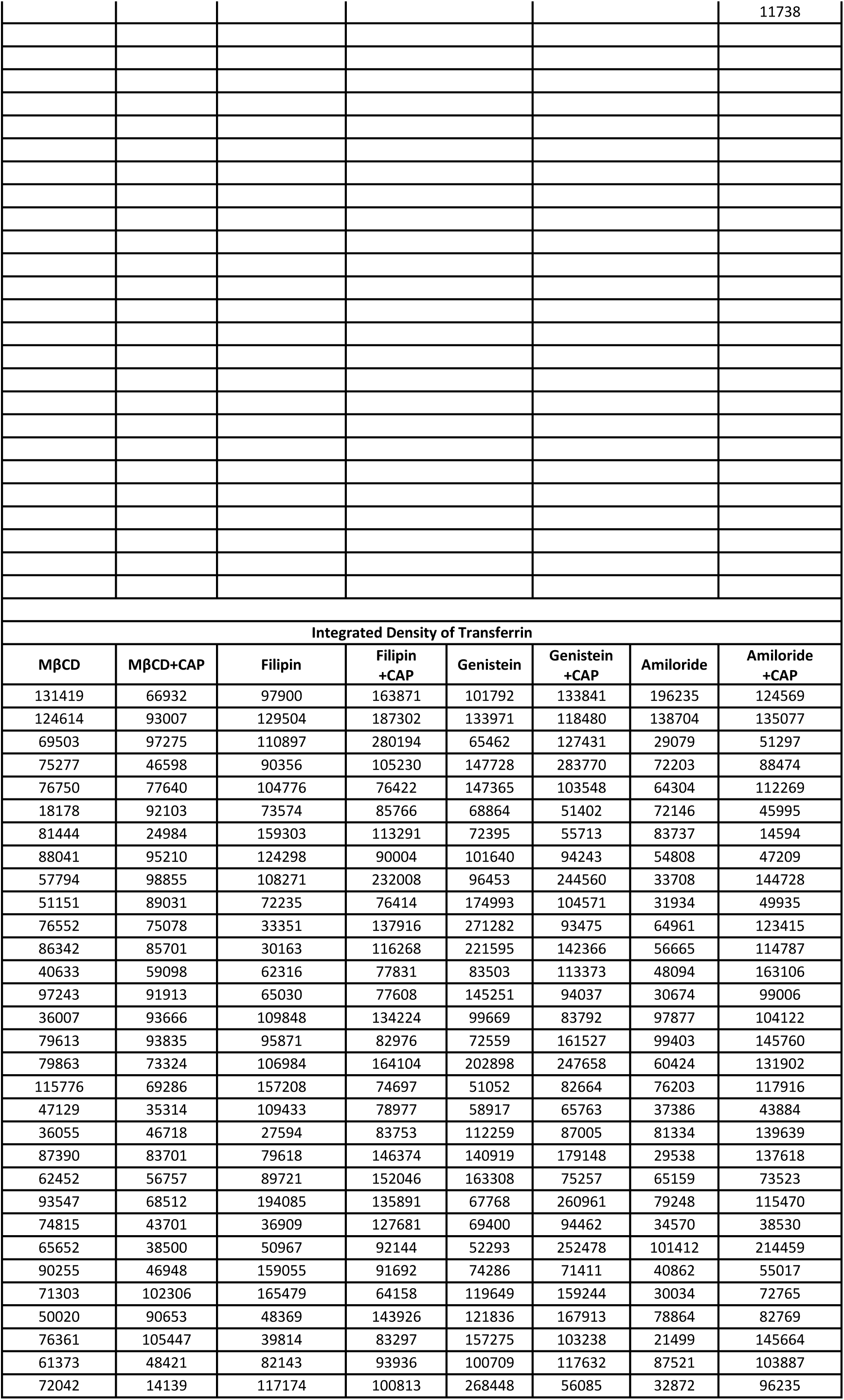
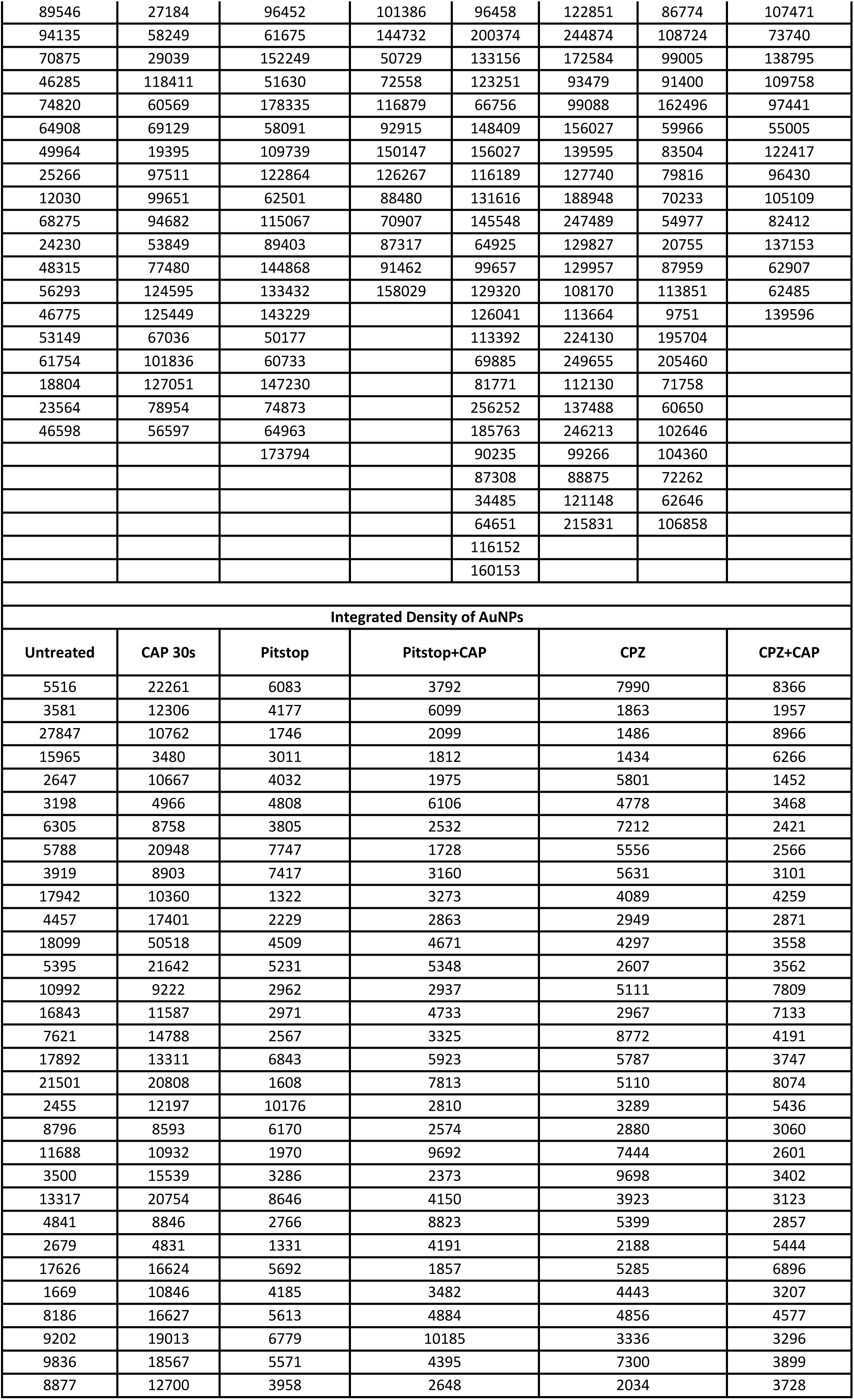
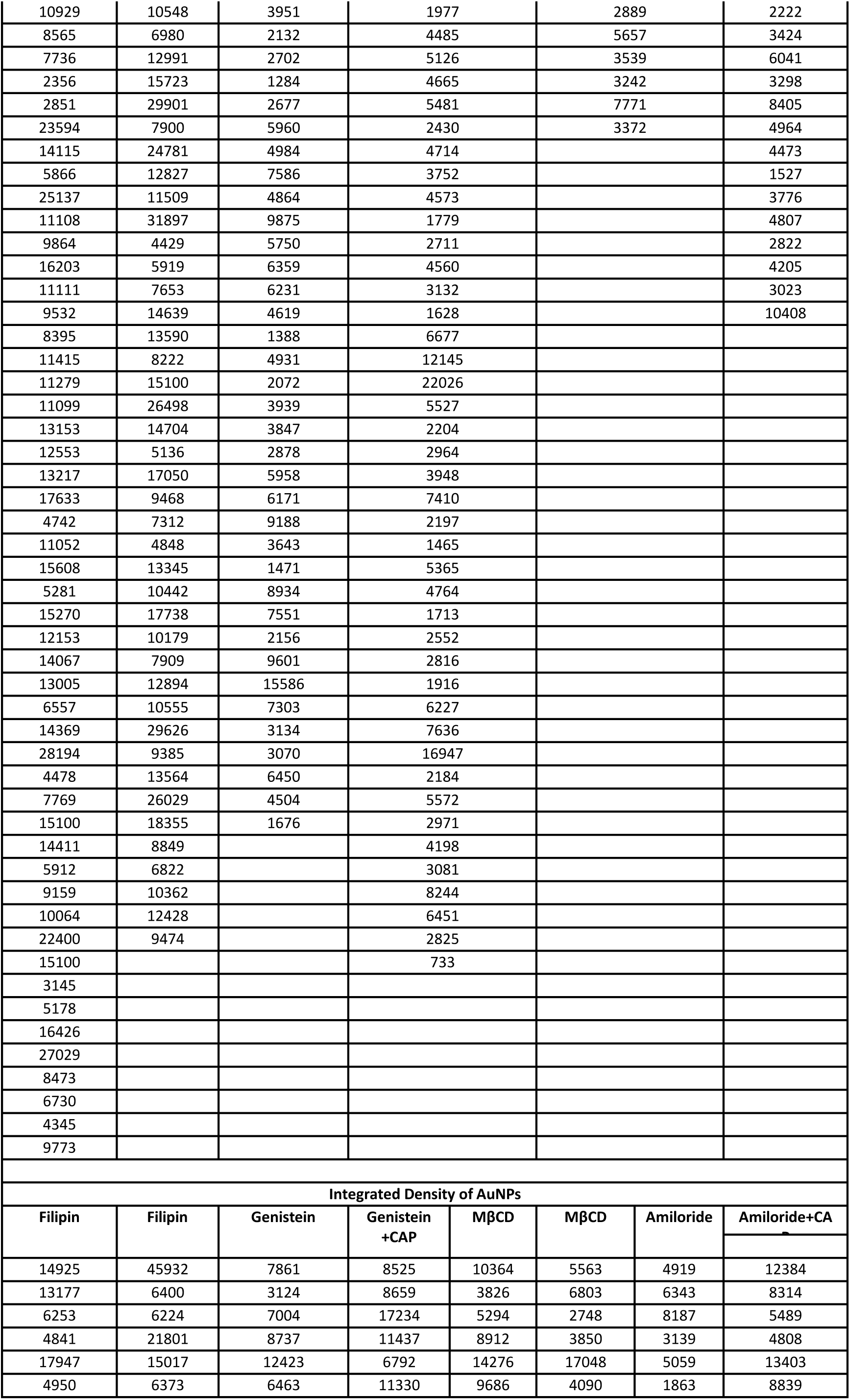
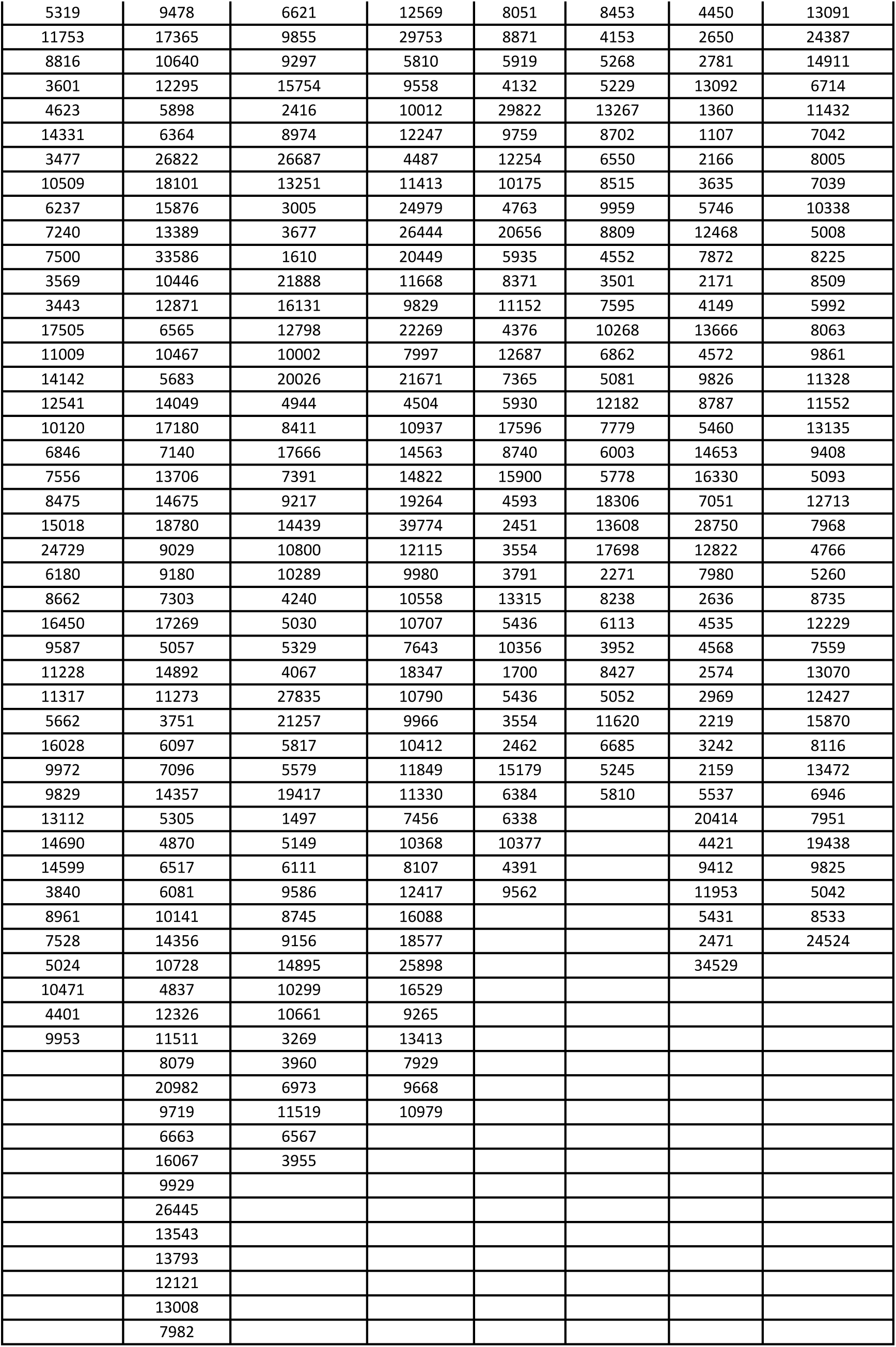

**Supplementary Table S4.**

| Integrated Density of AuNPs |
| --- |

| CAP30s Control | CAP 0s | CAP 0s | CAP 30s |
| --- | --- | --- | --- |
| 540329 | 172547 | 71657 | 81386 |
| 317286 | 431535 | 52885 | 26174 |
| 366683 | 78334 | 50113 | 83853 |
| 118864 | 227289 | 98205 | 62198 |
| 389856 | 182028 | 58882 | 113274 |
| 260748 | 95261 | 213664 | 176140 |
| 255370 | 67797 | 32246 | 90214 |
| 143714 | 155367 | 84077 | 155663 |
| 254835 | 143994 | 18060 | 167904 |
| 241522 | 409060 | 130518 | 52593 |
| 173393 | 91173 | 82556 | 22219 |
| 437882 | 98606 | 20741 | 19181 |
| 301708 | 372871 | 42321 | 16830 |
| 283998 | 42174 | 50000 | 36912 |
| 529897 | 192966 | 22797 | 11897 |
| 132739 | 43429 | 32307 | 67489 |
| 325045 | 171486 | 111229 | 22481 |
| 256487 | 155863 | 165385 | 45392 |
| 225093 | 218957 | 243576 | 12669 |
| 459960 | 332020 | 115717 | 14759 |
| 242832 | 163495 | 32769 | 59857 |
| 102314 | 178575 | 134059 | 29691 |
| 274004 | 43099 | 91085 | 36894 |
| 319820 | 116779 | 96918 | 46448 |
| 144878 | 33679 | 42816 | 24724 |
| 313094 | 281964 | 92220 | 41174 |
| 215681 | 474462 | 163157 | 12320 |
| 426863 | 78871 | 107430 | 42693 |
| 169499 | 25139 | 78092 | 54199 |
| 449778 | 157231 | 44965 | 26720 |
| 207286 | 25451 | 46103 | 68806 |
| 600642 | 105195 | 99071 | 50186 |
| 191247 | 124412 | 169342 | 18736 |
| 175692 | 128267 | 80229 | 16030 |
| 167735 | 312896 | 36048 | 55855 |
| 195195 | 129867 | 76657 | 106951 |
| 195351 | 127963 | 117916 | 17979 |
| 131726 | 352128 | 22666 | 39283 |
| 328548 | 257630 | 66817 | 17452 |
| 212497 | 116701 | 53062 | 34551 |
| 453971 | 134485 | 35270 | 10871 |
| 318022 | 113204 | 33485 | 43176 |
| 335974 | 203686 | 37665 | 6543 |
| 135597 | 81301 | 39651 | 20391 |
| 226104 | 258075 | 18179 | 9906 |
| 136796 | 117997 | 56467 | 43154 |
| 168061 | 189202 | 26238 | 33298 |
| 530325 | 354143 | 16544 | 6978 |
| 310895 | 179360 | 42583 | 10313 |
| 156015 | 249807 | 10222 | 7391 |
| 118700 | 104282 | 19974 | 18762 |
| 122974 | 189066 | 53493 | 56664 |
| 133195 | 84785 | 27174 | 8288 |
| 267536 | 141462 | 21953 | 72528 |
| 200762 | 345977 | 148609 | 24130 |
| 299249 | 77114 | 73347 | 13301 |
| 94514 | 47361 | 62139 | 18333 |
| 293151 | 43665 | 54334 | 20520 |
| 176776 | 9389 | 4625 | 17705 |
|  | 270307 | 6653 | 38519 |
|  | 31190 | 46708 | 28687 |
|  | 257884 | 50733 | 15938 |
|  | 155736 | 49224 |  |
|  | 345060 | 17503 |  |
|  | 43585 | 15541 |  |
|  | 81386 | 3876 |  |
|  | 23622 |  |  |
|  | 75993 |  |  |
|  | 65180 |  |  |
|  | 568575 |  |  |
|  | 318601 |  |  |
|  | 174400 |  |  |
|  | 52373 |  |  |
|  | 23718 |  |  |
|  | 59279 |  |  |
|  | 38939 |  |  |
|  | 75419 |  |  |
|  | 98606 |  |  |
|  | 24077 |  |  |
|  | 75776 |  |  |
|  | 41710 |  |  |
|  | 22592 |  |  |
|  | 70125 |  |  |
|  | 45832 |  |  |
|  | 98998 |  |  |
|  | 114681 |  |  |
|  | 126842 |  |  |
|  | 72896 |  |  |
|  | 161663 |  |  |

